# Microglial MorphOMICs unravel region- and sex-dependent morphological phenotypes from postnatal development to degeneration

**DOI:** 10.1101/2021.11.30.470610

**Authors:** Gloria Colombo, Ryan John A. Cubero, Lida Kanari, Alessandro Venturino, Rouven Schulz, Martina Scolamiero, Jens Agerberg, Hansruedi Mathys, Li-Huei Tsai, Wojciech Chachólski, Kathryn Hess, Sandra Siegert

## Abstract

Microglia contribute to tissue homeostasis in physiological conditions with environmental cues influencing their ever-changing morphology. Strategies to identify these changes usually involve user-selected morphometric features, which, however, have proved ineffective in establishing a spectrum of context-dependent morphological phenotypes. Here, we have developed MorphOMICs, a topological data analysis approach to overcome feature-selection-based biases and biological variability. We extracted a spatially heterogeneous and sexually-dimorphic morphological phenotype for seven adult brain regions, with ovariectomized females forming their own distinct cluster. This sex-specific phenotype declines with maturation but increases over the disease trajectories in two neurodegeneration models, 5xFAD and CK-p25. Females show an earlier morphological shift in the immediately-affected brain regions. Finally, we demonstrate that both the primary- and the short terminal processes provide distinct insights to morphological phenotypes. MorphOMICs maps microglial morphology into a spectrum of cue-dependent phenotypes in a minimally-biased and semi-automatic way.

## Introduction

Morphological characterization of neuronal shapes has provided important insights into the diversity of cell types related to their genetic and functional features (Gouwens et al., 2019; Jiang et al., 2015; Moschovakis et al., 1988). Numerous studies have tried to apply a similar morphological analysis on microglia (Adeluyi et al., 2019; Heindl et al., 2018; Kongsui et al., 2014; Morrison and Filosa, 2013; Morrison et al., 2017; Young and Morrison, 2018). Although they indicated a microglia heterogeneity (Bachstetter et al., 2015; Lawson et al., 1990; Stratoulias et al., 2019; Tan et al., 2019), no study has found a way to capture context-specific responses during development and degeneration. Moreover, the majority of these analysis rely on restricted microglia sample sizes underestimating their full morphological phenotypical spectrum. Detecting subtle changes in the microglial morphology along the spectrum would offer an early readout of their immediate responses to local environmental cues (Lawson et al., 1990), as microglia are sensitive to changes in neuronal activity (González-Scarano and Baltuch, 1999; Greter and Merad, 2013; Paolicelli et al., 2011; Pont-Lezica et al., 2011; Venturino et al., 2021).

Microglia morphological phenotype is commonly determined with user-selected features from a three-dimensional (3D) reconstructed branching tree: these features can include total process length, branch number, or number of terminal points. These scalar morphometric descriptors are then statistically compared across conditions. The drawback of this approach is the number and the type of selected features, which biases the biological readout: while too few selected features underrepresent a phenotypic difference, too many cause overfitting and introduce noise (Carlsson, 2009). Moreover, in contrast to neuronal morphological trees that are static on the gross structure, microglia processes are highly dynamic (Davalos et al., 2008; Nimmerjahn et al., 2005; Tremblay et al., 2011; Wu et al., 2015) as they constantly survey their local environment (Davalos et al., 2008). This introduces considerable intrinsic variability within the traced microglia population of a defined condition as well as the risk of selection bias to the extreme phenotypes. Establishing a reliable brain-region-specific morphological phenotype is critical for characterizing baseline morphology and tracking changes as deviations from the baseline.

To capture morphological phenotypes, complex morphological trees must be simplified with minimal information loss, retaining as many features as possible. Algebraic topology provides new strategies for solving this problem, as it focuses on the shape properties of geometric objects without the need of morphometrics (Kanari et al., 2017; Li et al., 2017). In particular, the topological morphology descriptor (TMD), which assigns a barcode to any three-dimensional tree, has been successfully applied for classification of cortical neuron morphologies (Kanari et al., 2017, 2020). When we first applied the TMD to ∼10,000 traced microglia across the rostro-caudal axis of seven selected adult brain regions, the data indicated a regional phenotype, but the diversity of the individual microglia obscured any well-defined separation. We therefore developed our MorphOMICs pipeline to overcome the major limitations of feature-selection-based analysis and biological data variability. MorphOMICs combines TMD with bootstrapping, dimensionality reduction, and data visualization techniques, enabling minimally-biased identification of the baseline phenotype. We found that adult microglia are brain-region-specific, with sex as a confounding factor. Remarkably, microglia from ovariectomized females did not match the corresponding brain regions of males nor females, underscoring the impact of sex on the morphological phenotypic spectrum. This was further strengthened when we analyzed postnatal development and degeneration. We found an early microglial sexual dimorphism, which gradually declined towards adulthood. In contrast, the sex-specific phenotype diverged during disease progression in two distinct Alzheimer’s disease models, 5xFAD and CK-p25, where females differ in their context-dependent response from males. When we aligned the trajectories of development and degeneration, we observed a phenotypic spectrum supporting that microglia morphology is context- and sex-dependent. Finally, we found that both short and long persistence bars are required to detect subtle changes within microglia morphology. MorphOMICs provides a minimally-biased, semi-automatic strategy for mapping a spectrum of microglia phenotypes across conditions.

## Results

### MorphOMICs uncovers adult microglial heterogeneity beyond intrinsic variability

To address how morphological phenotypes differ between microglia across brain regions, we immunostained the adult C57BL/6J mouse brain with the allograft inflammatory factor 1 (Iba1/Aif1) (Ito et al., 1998) for both sexes with at least biological triplicates. Then, we traced 9,997 cells and generated three-dimensional (3D) microglial skeletons from seven brain regions chosen to span the rostro-caudal axis with a preference for regions that are known to be affected in Alzheimer disease (Braak et al., 1989; Brar et al., 2009; Burns et al., 2005; DeKosky and Scheff, 1990; Gosche et al., 2002; Jacobs et al., 2018; Kazee et al., 1995; Kovács et al., 1999; Larner, 1997; Lehtovirta et al., 1995; Leuba et al., 2008; Ohm and Braak, 1987, 1989; Rao et al., 2012; Rombaux et al., 2010; Sinha et al., 1993; Stephen et al., 2010; Thompson et al., 2004; Wiesman et al., 2021; Zarow et al., 2003): the olfactory bulb (OB_mg_), frontal cortex (FC_mg_), dentate gyrus of the hippocampus (DG_mg_), primary somatosensory cortex (S1_mg_), substantia nigra (SN_mg_), cochlear nucleus (CN_mg_), and cerebellum (CB_mg_, **Fig. 1A**). When we utilized morphometrics that are commonly used in the field of microglial morphology (Kozlowski and Weimer, 2012; Morrison and Filosa, 2013; Zusso et al., 2012), we found non-significant differences across these brain regions with the exception of (CB and CN)_mg_ (**Supp. Fig. 1A**). We therefore applied the topological morphology descriptor (TMD) (Kanari et al., 2017; Li et al., 2017) for which each 3D skeleton was represented as a rooted tree with the microglial soma, processes, branching points, and process terminals (**Fig. 1B**, i). The TMD converts the tree as a persistence barcode, where each bar represents the persistent process lifetime in terms of the radial distance from the soma (Carlsson, 2009; Kanari et al., 2017). Every bar is then collapsed into a single point in the persistence diagram summarizing the process’s lifetime, which is then converted into a persistence image using Gaussian kernels (Adams et al., 2017) (**Fig. 1B**, ii, iii). The branching complexity is spatially represented by process length proportional to the distance from the diagonal (**Fig. 1B**, iv). To quantify the differences between microglial morphologies across brain regions, we computed the pairwise TMD distance between the average persistence images (Kanari et al., 2017). We identified clusters with (OB, FC, SN)_mg_ and (S1, DG)_mg_, whereas CN_mg_ and CB_mg_ were segregated (**Supp. Fig. 1B**). The average persistence images also did not differ strongly (**Fig. 1C**). When we looked at the individual persistence images, we found a wide variance that makes it challenging to distinguish regional phenotypes (**Supp. Fig. 1C**). We note that this dispersion is not driven by an animal-based batch effect (**Supp. Fig. 1D**).

**Figure 1.**
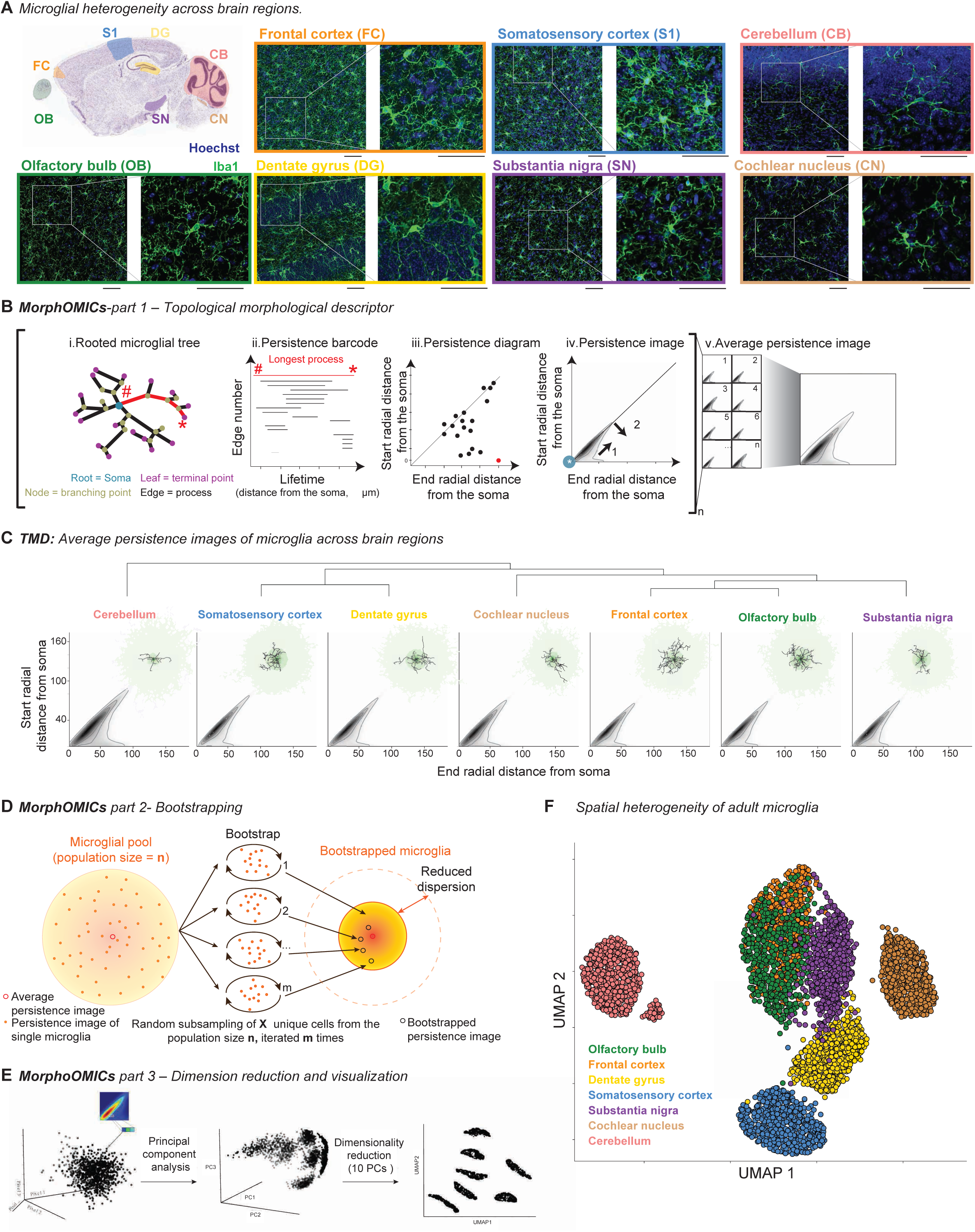
MorphOMICs dissects microglial morphology in adult healthy brains. **A:** Sagittal view of the murine brain (Image credit: Allen Institute) with annotated brain regions: olfactory bulb (OB), frontal cortex (FC), dentate gyrus (DG), somatosensory cortex (S1), substantia nigra (SN), cochlear nucleus (CN), and cerebellum (CB). Next, confocal images of immunostained microglia (Iba1, allograft inflammatory factor 1, green) and cell nuclei (Hoechst, blue) from adult C57BL/6J mice with zoom-in. Scale bar: 50 μm. **B:** Schematic of MorphOMICs pipeline covering topological morphology descriptor (TMD). Each traced microglia is converted into a rooted tree (i, start (#) and end (*) of longest process in red), and from there into a persistence barcode (ii), a persistence diagram (iii, with each bar collapsed to a point), and a persistence image (iv) with grey-scaled process density in space. Blue spot: soma location. Arrow 1 indicates distance from the soma and arrow 2 process length, which increases the further away from the diagonal. n persistence images are summarized to an average persistence image. **C:** Average persistence images of the seven analyzed brain regions organized by hierarchical clustering (see also **Supp. Fig. 1B**). Top corner: representative microglia. The darker the green, the higher the frequency distribution of the processes. **D:** Schematic of MorphOMICs pipeline covering bootstrapping. Microglial population (n) contains individual persistence images. In the center: average persistence image. x unique persistence images are drawn from each of n microglial pools to generate a bootstrapped persistence image. Repeating this process m times forms the bootstrapped pool. **E:** Schematic of MorphOMICs pipeline covering dimension reduction and data visualization with Uniform Manifold Approximation and Projection (UMAP). Left: each persistence image is pixelated, and each pixel represents a dimension. Middle: reducing dimensions with principal component (PC) analysis. Right: further dimensionality reduction based on the first ten PCs. **F:** UMAP plot of MorphOMICs-analyzed microglia, color-coded for each brain region. Each dot represents a bootstrapped persistence image (**D**). Data for each brain region from at least six animals, both sexes together.

To overcome this intrinsic variability within a microglial population, we developed MorphOMICs, which combines TMD with random subsampling of persistence images, dimensionality reduction, and data visualization strategies. Bootstrapping randomly draws, without replacement, a user-defined number of unique persistence images (x) from a microglial population pool (n) and iteratively generates bootstrapped persistence images (**Fig. 1D**). To display these bootstrapped persistence images for each brain region, we applied the nonlinear dimensionality reduction technique UMAP (Uniform Manifold Approximation and Projection, **Fig. 1E**), which converts the high-dimensional persistence images into a reduced 2D representation while preserving their global structure (Mcinnes et al., 2020). While local distances are preserved in UMAP compared to t-SNE (Van Der Maaten and Hinton, 2008), the point’s actual position in the reduced space is irrelevant. After controlling for the bootstrapped to microglial population pool size ratio (**Supp. Fig. 2A-E**, **Supplementary Text**), we applied MorphOMICs to our data. The UMAP plot exhibited a spatial separation similar to that of the distance matrix in **Supp. Fig. 1B**, with CB_mg_ separated from the other brain regions and (OB, FC, SN)_mg_ occupying a well-defined area in the UMAP space (**Fig. 1F**). Interestingly, MorphOMICs revealed that OB_mg_ and FC_mg_ are intermingled, while DG_mg_ and S1_mg_ formed distinct clusters. Importantly, these cluster segregations were stable even if we changed UMAP’s hyperparameters (**Supp. Fig. 2F**) or when we applied t-SNE visualization instead (**Supp. Fig. 2G**). More complex morphological relationships between brain regions can exist as exhibited by CN_mg_. When we represented the persistence barcodes with stable ranks instead of persistence images (Agerberg et al., 2021; Riihimäki and Chacholski, 2018), the region-specific phenotypes were maintained (**Supp. Fig. 2H**). Furthermore, stable ranks allowed us to discriminate microglia in to different brain regions with a classification accuracy that reflects the separation between brain regions in the UMAP space (**Supp. Fig. 3A**). An alternative morphological simplification that is commonly performed in the literature is Sholl analysis, which calculates the number of processes that intersect concentric spheres centered on the soma with a user-defined radius (Sholl, 1953). When we applied Sholl analysis we could not recapitulate entirely the spatial segregation captured by MorphOMICs (**Supp. Fig. 3B**). Even if we applied bootstrapping to Sholl-analysis, we could only dissect the regional heterogeneity for CB_mg_ and CN_mg_ (**Supp. Fig. 3C**). Interestingly, the clusters become less distinct by increasing the Sholl step size radius confirming the superiority of MorphOMICs. Overall, these data indicate that adult brain regions have well-defined microglia morphological phenotypes, which is uncovered by MorphOMICs.

### The microglial phenotype displays sexual dimorphism in a brain-region-dependent manner

The extent of microglial sexual dimorphism across brain regions is only partially understood (Han et al., 2021; Lenz et al., 2013; Nelson et al., 2017). We applied MorphOMICs to all brain regions, and compared males and females within the UMAP space (**Fig. 2A-B**). Each brain region occupied a unique cluster in the plot, where CB_mg_ and CN_mg_ were most divergent. Strikingly, most brain regions separated female and male microglia, with CB_mg_, CN_mg_, SN_mg_, S1_mg_, and OB_mg_ forming close but spatially separated clusters. In contrast, ♂/♀DG_mg_ and FC_mg_ highly overlapped, suggesting rather minor morphological differences between the sexes. Interestingly, compared to **Supp. Fig. 1B** the (FC, OB)_mg_ cluster broke up: ♂FC_mg_ and ♂OB_mg_ formed spatially separated clusters, whereas ♀(FC, OB)_mg_ were intermingled.

**Figure 2.**
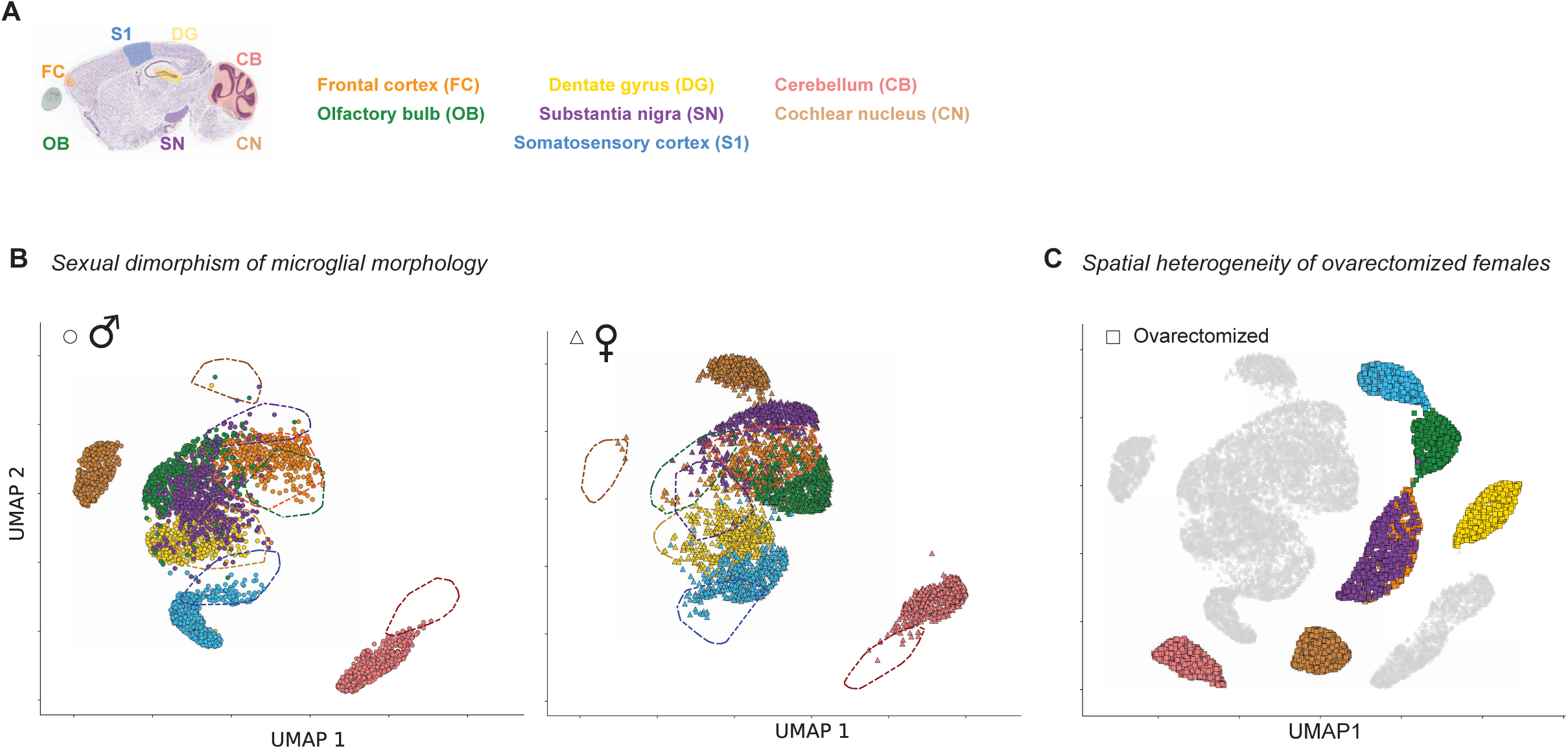
MorphOMICs identifies sexually-dimorphic microglial morphology in healthy adults. **A:** Sagittal view of analyzed brain regions: olfactory bulb (OB), frontal cortex (FC), dentate gyrus (DG), somatosensory cortex (S1), substantia nigra (SN), cochlear nucleus (CN), and cerebellum (CB) (Image credit: Allen Institute (Lein et al., 2006)). **B:** UMAP plot of MorphOMICs-analyzed microglia for each brain region color-coded for males (left) or females (right) with dashed lines *vice versa* as reference. Each dot represents a bootstrapped persistence image. CN_mg_: t = 3.504, df = 15.1, p-value = 0.00312. OB_mg_: t = 2.401, df = 16.864, p-value = 0.0282. CB _mg_: t = 1.2564, df = 17.327, p-value = 0.2257. FC _mg_: t = 1.6236, df = 16.275, p-value = 0.1237. SN _mg_: t = 1.6261, df = 12.901, p-value = 0.1281. DG _mg_: t = 0.68892, df = 19.669, p-value = 0.4989. S1_mg_ t = 1.5618, df = 17.518, p-value = 0.1362. n_♀_=11, n_♂_= 11. **C:** UMAP plot of MorphOMICs-analyzed microglia in ovarectomized females. Ovariectomized brain regions are highlighted and the non-ovarectomized counterpart is shown in grey as a reference (see also **Supp. Fig. 3B**).

The morphological differences could depend on the microglia distribution; therefore, we determined the microglia density for each brain region and sex. We found that the density was the lowest for both sexes in CB_mg_ (**Supp. Fig. 4A**) and only CN_mg_ and OB_mg_ showed a significant sexual dimorphism. This is reflected within the UMAP space, where both brain regions showed the strongest separation (**Fig. 2B**). To determine whether this sex-specific intermingling is hormone-dependent, we traced microglia from adult females that we ovariectomized at P20 (♀_ov,_ **Supp. Fig. 4B**) before they start the estrous cycle and enter puberty (Caligioni, 2009). We found that the ♀_ov_FC_mg_ cluster no longer intermingled with ♀_ov_OB_mg_ in ovariectomized females but instead fused with ♀_ov_SN_mg_ (**Fig. 2C**). This is surprising, as in non-ovariectomized mice, ♀SN_mg_ was close to but distinct from the intermingled ♀(FC, OB)_mg_. When we compared non-ovariectomized to ovariectomized females, we found that in the UMAP space ovariectomized females formed distinct clusters, spatially separated from their non-ovariectomized counterparts (**Fig. 2B-C**). These results demonstrate the existence of brain-region-specific, sexually dimorphic phenotype, and that interfering with estrogen production before puberty affects microglial heterogeneity.

### Sexual dimorphism affects microglial morphology during development

Microglia originate in the yolk sac and infiltrate the nervous system early during embryonic development (Ginhoux et al., 2013). After microglia occupy a brain region, their morphology gradually becomes more branched during postnatal neuronal circuit refinement (**Fig. 3A**) (Ben-Ari, 2002; Ginhoux et al., 2010; Kroon et al., 2019; Perez-Pouchoulen et al., 2015; Ruusuvuori et al., 2004; Valeeva et al., 2016; Wong et al., 2005; Yang et al., 2014). To determine whether microglial heterogeneity and dimorphic phenotype already exist within the first postnatal weeks and before the onset of puberty, we sampled microglia from all seven brain regions at postnatal days 7, 15, and 22 (**Supp. Fig. 5A-B**) and highlighted either each brain region or the developmental time point in the UMAP plots (**Fig. 3B-C**, respectively). In all seven brain regions, no postnatal time points overlapped with the adult microglia (**Fig. 3B**), reflecting their morphological spectrum during development. When we analyzed each time point individually, we found that at P7, all brain regions are distinct but occupy the same cluster, which shifted to a different cluster at P15 (**Fig. 3C**). Interestingly, CN_mg_ and DG_mg_ segregated and remained distinct from the other brain regions at P15 and P22, with CB_mg_ joining them at P22. Between P22 and adulthood, the clusters diverged to their adult microglial heterogeneity.

**Figure 3.**
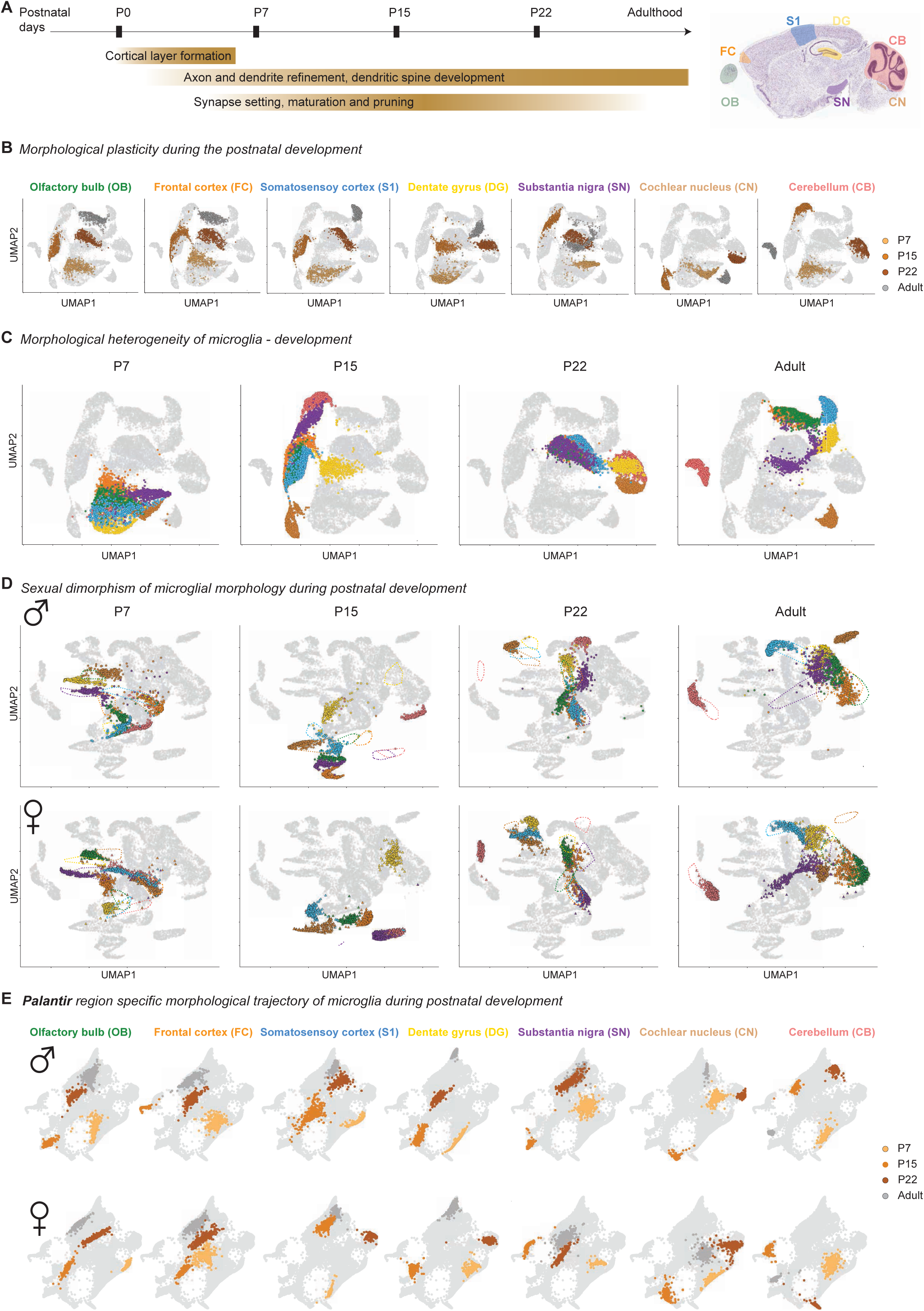
Microglial phenotypes during postnatal development. **A:** Left: schematic of postnatal brain development with key aspects of brain maturation and neuronal circuitry refinement. Right: sagittal view of analyzed brain regions: olfactory bulb (OB), frontal cortex (FC), dentate gyrus (DG), somatosensory cortex (S1), substantia nigra (SN), cochlear nucleus (CN) and cerebellum (CB). (Image credit: Allen Institute (Lein et al., 2006)). **B**-**D:** UMAP plots of MorphOMICs-analyzed microglia across seven brain regions in Cx3cr1-GFP^+/-^ mice (**B**) at P7 (light brown), P15 (orange), P22 (dark brown), and adults (dark grey); and color-coded brain regions for both sexes (**C**) and for each sex independently with dashed lines *vice versa* as reference (**D**) for each developmental time point. Each dot represents a bootstrapped persistence image. **E:** Palantir reconstruction of microglia morphological trajectory from (**D**) with highlighted P7 (light brown), P15 (orange), P22 (dark brown) and adults (dark grey) for each brain region.

Next, we investigated whether sexual dimorphism affects microglial phenotypic spectrum during development. To do this, we applied the MorphOMICs pipeline to males and females separately (**Fig. 3D**, **Supp. Fig. 5C**). Surprisingly, we found that the clusters shown in **Figure 3C** split, leading to well-defined male and female clusters for each brain region at P7. With brain maturation, ♀/♂ clusters in S1_mg_, DG_mg_, and FC_mg_ converged, while those in OB_mg_, SN_mg_, CB_mg_, and CN_mg_ remained distinct. To follow this sexual dimorphism along the developmental trajectory, we ordered the bootstrapped persistence images with the Palantir algorithm (Setty et al., 2019), which uses principles from graph theory and Markov processes to infer a pseudo-temporal trajectory (**Fig. 3E**). In the Palantir space, nearby points indicate similar persistence images, thereby assuming a gradual transition in their morphologies and the continuous sequence of points define a trajectory. The developmental trajectories were similar between brain regions, with P7 and P22 clusters being the furthest from and the closest to the adult, respectively. In contrast, P15 shifted laterally from the P7-P22 trajectory and occupied the outermost position in nearly all the brain regions, indicating a unique microglial context-dependent response that coincides with neuronal circuit synapse refinement (Kroon et al., 2019; Wong et al., 2005; Yang et al., 2014).

### The correlation between microglial phenotypic spectrum and reactivity is region specific in 5xFAD

Synaptic loss combined with amyloid plaque deposition are common signs of Alzheimer’s disease, with the neocortex and the hippocampus being the most affected brain regions (Serrano-Pozo et al., 2011; Terry, 2000). Microglial morphology alters during the progression of Alzheimer’s disease (Hemonnot et al., 2019; Taipa et al., 2018) but the disease phenotype of microglia in directly- and indirectly-affected brain regions, as well as the impact on the sexual dimorphism, is not well understood (Congdon, 2018; Gamache et al., 2020; Manji et al., 2019). To address this with MorphOMICs, we traced microglial morphologies in the 5xFAD mouse model, which recapitulates a familial form of Alzheimer’s disease (**Fig. 4A**, **Supp. Fig. 6A**) (Oakley et al., 2006). We traced morphologies for all seven brain regions and focused on animals that were three and six months old (5xFAD_3m_ and 5xFAD_6m_, respectively, **Supp. Fig. 6B**) because amyloid plaques occur first in the deep cortical layers at three months, followed by the hippocampus, coinciding with spine loss and memory deficits around 6 months (Oakley et al., 2006). As anticipated, microglia in the 5xFAD_3m_ group exhibited a disease phenotype in which all brain regions were distinguishable from controls. The 5xFAD_6m_ group formed a “disease-associated cluster” in the UMAP space, with the exception of CB_mg_ (**Fig. 4B**). FC_mg_, S1_mg_, and DG_mg_ in 5xFAD_3m_ mice already occupied this disease-associated cluster. By subtracting representative bootstrapped persistence images, we observed that microglia from 5xFAD_6m_ have more short and long persistent processes close to the soma compared to controls (**Supp. Fig. 6C**). Moreover, the 5xFAD_6m_ - control TMD distances reflect the relative phenotypic changes in the UMAP space in **Fig. 4B**.

**Figure 4.**
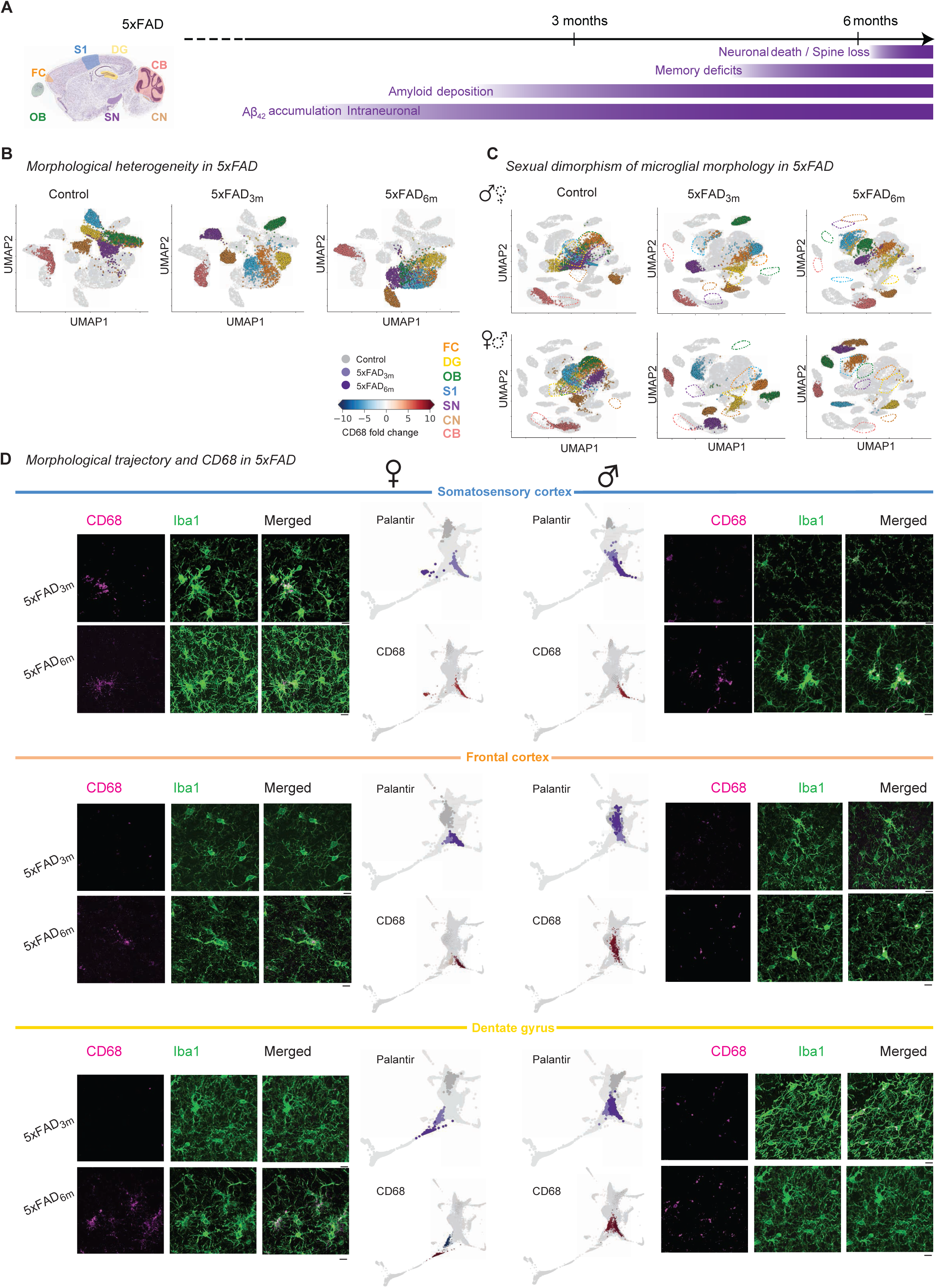
Microglia phenotypic spectrum in 5xFAD transgenic model of neurodegeneration is sexually dimorphic. **A:** Left: sagittal view of analyzed brain regions: olfactory bulb (OB), frontal cortex (FC), dentate gyrus (DG), somatosensory cortex (S1), substantia nigra (SN), cochlear nucleus (CN) and cerebellum (CB). (Image credit: Allen Institute (Lein et al., 2006)). Right: timeline of degeneration events in 5xFAD transgenic mouse model. **B**-**C:** UMAP plots of MorphOMICs- analyzed microglia across seven brain regions for the following conditions: control, 5xFAD_3m_ (3 months), and 5xFAD_6m_ (6 months) with both sexes (**B**) or for each sex separated (**C**). Each degeneration time point is highlighted in a separate UMAP. Each dot represents a bootstrapped persistence image. **D:** Representative confocal images of microglia (Iba1, green) and lysosome (CD68, magenta), followed by Palantir reconstruction of microglial trajectory with corresponding color-coded CD68 fold change below from females (left side) and males (right side) for control, 5xFAD_3m_, and 5xFAD_6m_ in S1, FC, and DG. Scale bar: 10 μm. Fold change < 0 blue; > 0 red.

When we included sex in our MorphOMICs analysis, the disease cluster was less pronounced (**Fig. 4C**). Instead, microglia demonstrated higher morphological heterogeneity in 5xFAD_3m_ compared to control, with males and females occupying distinct clusters. When we applied Palantir to identify sex-dependent disease trajectories, we observed sexual dimorphism (**Fig. 4D**, **Supp. Fig. 7A**), especially in one of the first affected brain region, S1. ♀S1_mg_ seem to precede ♂S1_mg_: ♀S1_mg_ clusters already overlapped in 5xFAD_3m_, whereas those in males first overlapped at 6 months (**Fig. 4D**). Such a difference in overlap was less obvious for FC_mg_ and DG_mg_, which is likely influenced by the limited number of selected time points over the course of the pathology. On the other hand, both regions display a phenotypic spectrum along the disease trajectory. To link microglial phenotype to their reactivity, we performed immunostaining for the endosomal-lysosomal marker CD68 (Chistiakov et al., 2017). We then computed the fold-change compared to the control CD68 volume within Iba1^+^-cells and overlaid the CD68 fold-change on the Palantir trajectory (**Fig. 4D**). In ♀S1_mg_, CD68 increased already at 3 months while this only occurred in ♂S1_mg_ at 6 months, confirming that the shift along the morphological spectrum happens earlier in females. For the other brain regions, this effect was less obvious. In FC, ♂FC_mg_ appeared to have a reactive microglial state at 3 months based on the CD68 staining, however their morphology overlapped with the control. In contrast, ♀FC_mg_, 5xFAD_3m_ phenotype segregated from the control, although the CD68 was only upregulated in 5xFAD_6m_. In the DG, ♀DG_mg_ moved along the disease trajectory faster than ♂DG_mg_, as for S1_mg_, while the CD68 response in DG_mg_ is closer to that of FC_mg_. We also applied Palantir trajectories to the other brain regions, since plaques deposition has been reported in the olfactory bulb and brainstem (Struble and Clark, 1992). We found a strong sexual dimorphism in microglial morphology in these brain regions, with less-obvious trajectory changes (**Supp. Fig. 7C**). CB_mg_ was the only exception, remaining mainly unaffected in 5xFAD mice, which is consistent with previous literature (Oakley et al., 2006). Overall, the 5xFAD data indicate that the link between microglial disease phenotype and reactivity state depends on the brain region.

### Female microglia exhibit an earlier shift along the morphological spectrum in a model of sporadic neurodegeneration

An alternative model with faster onset and disease progression is the CK-p25 model for sporadic Alzheimer-like degeneration (Cruz and Tsai, 2004; Cruz et al., 2003; Fischer et al., 2005). Upon doxycycline withdrawal, p25 expression is induced in CamKII^+^ forebrain neurons, resulting in neurotoxic activity of the cyclin-dependent kinase Cdk6 (Camins et al., 2006). Within two weeks, CK-p25 mice develop progressive neuronal and synaptic loss, forebrain atrophy, aberrant amyloid-precursor protein processing, hyper-phosphorylation of tau (Cruz et al., 2003, 2006), and at later stages neurofibrillary tangle-like pathology (Cruz et al., 2003, 2006) (**Fig. 5A**). We reconstructed microglial morphologies from CK-p25 mice at 1, 2, and 6 weeks (CK-p25_1w_, CK-p25_2w_, CK-p25_6w_, respectively, **Supp. Fig. 8A-B**) and applied MorphOMICs. Similar to 5xFAD, all seven brain regions started to segregate from the control at 1 week and occupied a disease-associated cluster in CK-p25_6w_, with CB_mg_ and CN_mg_ staying distinct (**Fig. 5B**). In particular, FC_mg_ reached this cluster already at 2 weeks, while OB_mg_, DG_mg_, S1_mg_, and SN_mg_ only at 6 weeks. As in 5xFAD, we confirmed that microglia during CK-p25 have more short and long processes close to the soma compared to controls (**Supp. Fig. 8C**), and that the CK-p25_6w_ - control TMD distances reflect the relative phenotypic changes in the UMAP space in **Fig. 5B**.

**Figure 5.**
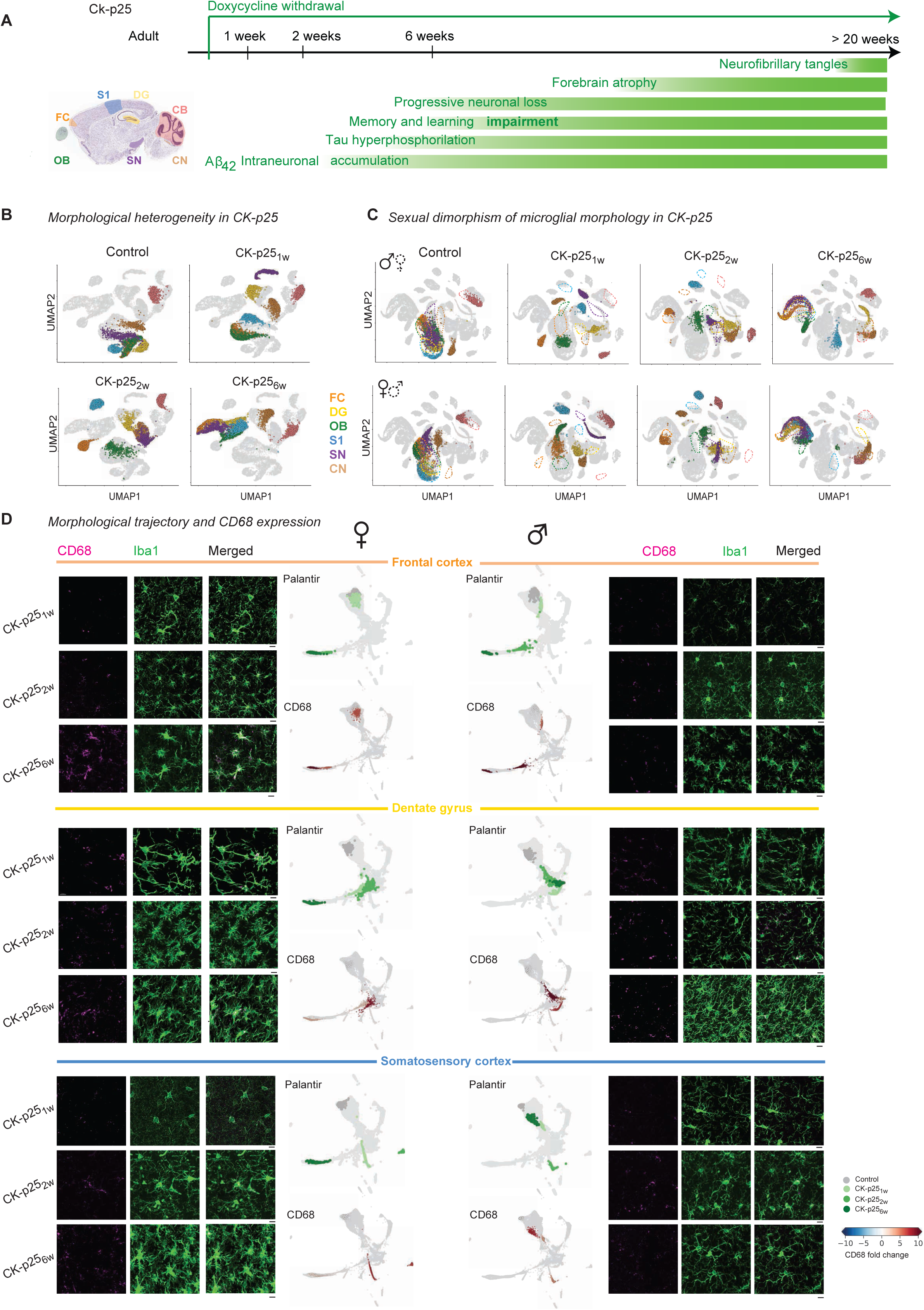
The microglia phenotype of females in CK-p25 model of neurodegeneration exhibit an earlier morphological shift than in males. **A:** Left: sagittal view of analyzed brain regions: olfactory bulb (OB), frontal cortex (FC), dentate gyrus (DG), somatosensory cortex (S1), substantia nigra (SN), cochlear nucleus (CN) and cerebellum (CB). (Image credit: Allen Institute (Lein et al., 2006)). Right: timeline reporting neurodegeneration events upon doxycycline withdrawal in the CK-p25 transgenic mouse model. **B**-**C:** UMAP plots displaying microglial morphological heterogeneity in adult control mice and CK-p25 mice 1, 2 and 6 weeks after doxycycline withdrawal across all the analyzed brain regions for both sexes (**B**) or for each sex separated (**C**). Each dot represents a bootstrapped persistence image, and each UMAP highlights a distinct degeneration time point. **D:** Representative confocal images of microglia (Iba1, green) and lysosome (CD68, magenta) from females (left) and males (right) in CK-p25 mice 1, 2, and 6 weeks after doxycycline withdrawal in S1, FC, and DG. Scale bar: 10 μm. Palantir reconstruction of microglial morphological trajectory and corresponding color-coded CD68 fold change of female (central column, left) and male (central column, right). Fold change < 0 blue; > 0 red.

When we applied MorphOMICs to each sex individually, we found that ♀(SN, FC, OB, DG, and S1)_mg_ reached the disease cluster at 6 weeks, while in males, ♂DG_mg_ and ♂S1_mg_ stayed distinct (**Fig. 5C**). Palantir replicated the UMAP and displayed an arm along the trajectory, on which microglial morphology from later disease stages accumulated (**Supp. Fig. 9A**). We note that neither CN_mg_ nor CB_mg_ reached this disease-associated arm as expected, due to the low expression of CamKII in these brain regions (Wang et al., 2013). Comparison of the sex-specific Palantir projections also showed that ♀FC_mg_ preceded ♂FC_mg_ in CK-p25_2w_ (**Fig. 5D**). We could replicate the same dynamics with Monocle, an alternative algorithm which uses reverse graph embedding to infer a pseudo-time trajectory (**Supp. Fig. 9B**) (Trapnell et al., 2014).

When we overlaid the CD68 fold-change compared to control adults over the Palantir FC_mg_ trajectory, we found that the CD68 fold-change gradually increased in ♂FC_mg_, while ♀FC_mg_ in CK-p25_6w_ showed a sudden increase (**Fig. 5D**). We also observed that the morphology did not correspond to CD68 in ♀DG _mg_ and ♀S1_mg_ at 6 weeks: the morphology reached the disease-associated arm but did not show any increased CD68 fold change. Neither ♂DG_mg_ nor ♂S1_mg_ reached the disease-associated arm, which we also confirmed with Monocle (**Supp. Fig. 9B**). On the other hand, ♂DG_mg_ and ♀DG_mg_ showed their highest CD68 fold change at 2 weeks and occupied a similar cluster in the Palantir space (**Fig. 5D**). This suggests that the microglial response might be associated with the transient effect of p25 expression, which has been shown to enhance long-term potentiation and improve hippocampus-dependent memory, before inducing neurodegeneration, gliosis, and severe cognitive decline at 6 weeks (Fischer et al., 2005). For those brain regions that were less affected, dimorphic microglial phenotype was less pronounced (**Supp. Fig. 9C**). In both sexes, SN_mg_ and OB_mg_ in CK-p25_6w_ reached the disease-associated arm, whereas in CB_mg_ and CN_mg_, neither sex nor disease progression influenced morphology (**Supp. Fig. 9D**). Overall, the CK-p25 model exhibited strong dimorphic phenotype spectrum in favor of females, which precede their male counterparts in a brain-region-specific manner.

### Microglial phenotype during early developmental integrates into the disease spectrum

Until now, we have treated microglial morphology separately for development and disease. Since both the development and the disease phenotypes result from a shift along the morphological spectrum, we were interested in how these conditions integrate along the pseudo-temporal trajectory. To achieve this, we performed MorphOMICs for all brain regions, including developmental time point P7, 5xFAD, and CK-p25 for each sex separately and extracted the trajectory with Palantir. We first focused on the female trajectory for CK- p25_6w_ in ♀FC_mg_ and ♀DG_mg_ identifying the control-to-disease spectrum (**Fig. 6A**). In ♀FC_mg_, the P7, 5xFAD_3m_, and 5xFAD_6m_ groups formed a continuum with the latter reaching out towards the CK-p25_2w_ group. Interestingly, ♀DG_mg_ mimicked a similar trajectory but with both 5xFAD and early CK-p25 forming a cluster distant from the control, and P7 group reaching out towards CK-p25_6w_. All ♀CB_mg_, which is the less affected region during neurodegeneration, segregate in the Palantir space, excluding P7.

**Figure 6.**
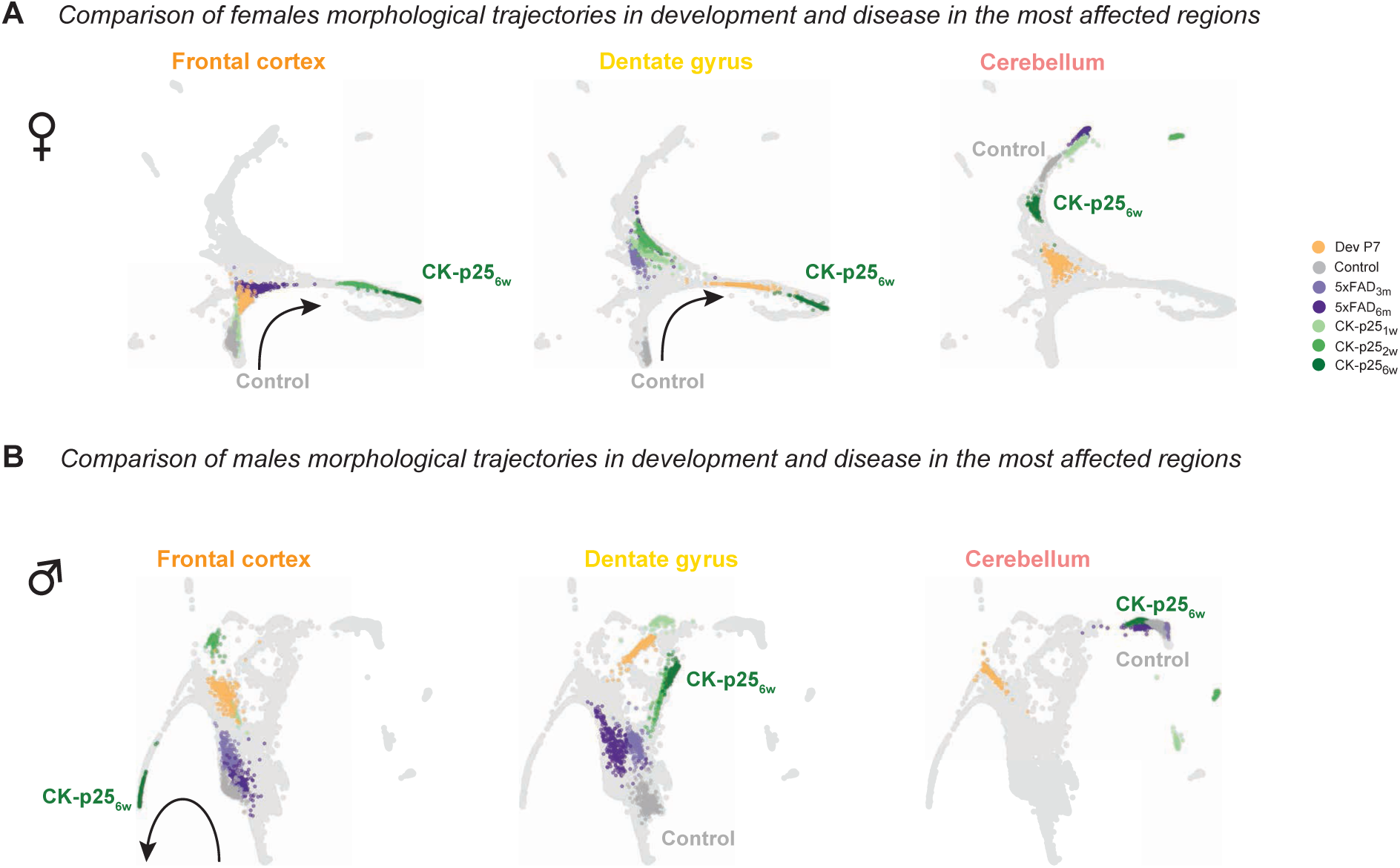
Early developmental phenotype integrates into the disease spectrum. **A**-**B:** Palantir reconstructions of microglial trajectory in females (**A**) and males (**B**) for FC, DG, and CB in control (grey), P7 (light brown), CK-p25_1w_ (1 week, light green), CK-p25_2w_ (2 weeks, green), CK-p25_6w_ (6 weeks, dark green), 5xFAD_3m_ (3 months, light purple), and 5xFAD_6m_ (6 months, purple) conditions. Each brain region is highlighted in a separate Palantir plot. Black arrow: control-to-disease spectrum.

In males, disease phenotypes evolved more slowly than in females, with only ♂FC_mg_ reaching the disease-associated arm at CK-p25_6w_ (**Fig. 6B**). Like in females, we observed both 5xFAD groups close to the control, followed by intermingled P7, CK-p25_1w_ and, subsequently, CK-p25_2w_. ♂DG_mg_ displayed similar phenotypic spectrum compared to ♂FC_mg_ for both 5xFAD groups, shifting towards the CK-p25_2w_-CK-p25_6w_ cluster. ♂CB_mg_ behaved similarly to females in the 5xFAD model (**Fig. 4C**). When we looked at the indirectly affected brain regions, the control-to-disease spectrum was less prominent (**Supp. Fig. 10A-B**). Overall, our data show that microglia display a spectrum of phenotypes, with P7 occupying distinct parts of the trajectory in a brain-region-dependent manner. This suggests that the microglial phenotype is comparable between normal development and disease, which are both environments with neuronal circuit remodeling.

### MorphOMICs maximizes the extraction of morphological information

MorphOMICs enabled us to establish both an adult sexual dimorphic phenotype and a morphological spectrum during development and degeneration for each brain region. Such a reconstruction is not straightforward with common morphological feature selection (**Supp. Fig. 1A**). We could not replicate the sexually dimorphic control-to-disease spectrum from **Fig. 5D** on microglia from females and males in CK-p25 mice with statistical comparisons of process length, number of branches, and terminal, and branching points (**Supp. Fig. 11A**). We also lost information of the morphological spectrum when we applied bootstrapping approaches to the extended set of non-interdependent morphometric quantities on the CK-p25 FC_mg_ (**Fig. 7A**). The same loss was observed applying the bootstrap method to microglia in the 5xFAD model and during development (**Supp. Fig. 11B-C**). This suggests that MorphOMICs preserves certain intrinsic properties of the reconstructed tree after dimensionality reduction.

**Figure 7.**
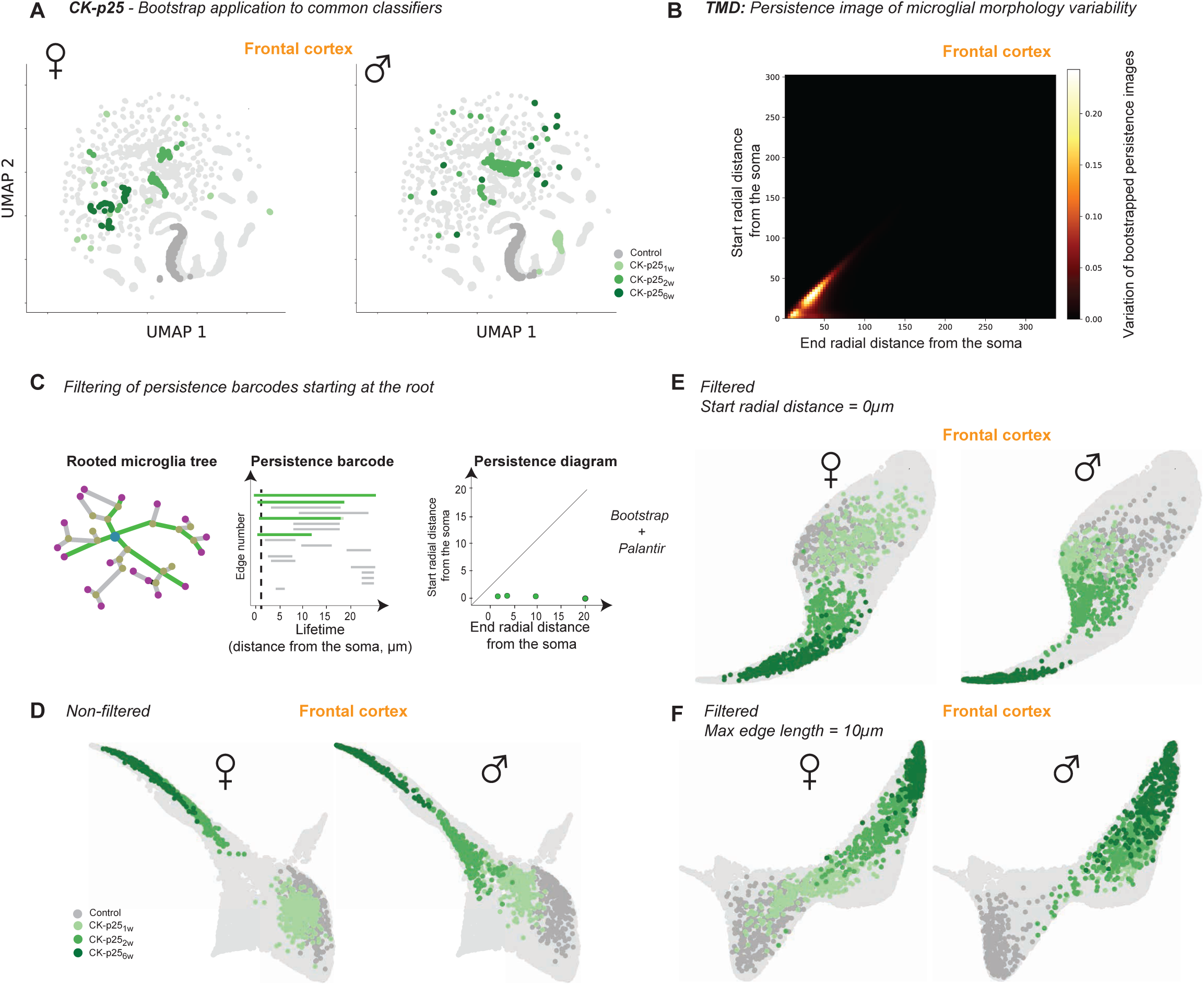
MorphOMICs applied to primary processes reiterates sexual dimorphism in CK-p25 mice. **A:** Left: box plots for the selected features dendritic length, number of branches, and terminal and branching points of control (n_♀_= 926, n_♂_= 894), and CK-p25_1w_ (n_♀_= 219, n_♂_= 194), CK- p25_2w_ (n_♀_= 264, n_♂_= 492), CK-p25_6w_ (n_♀_= 858, n_♂_= 462) mice (1-, 2- and 6-weeks after doxycycline withdrawal, respectively) in the frontal cortex (FC). Right: matrices showing color-coded p-values for the pairwise comparison of each morphometric **A**: UMAP representations based on multiple classical morphometrics (see **Supp. Table 4**) during CK- p25_6w_ degeneration in females (left) and males (right). Highlighted clusters are FC for control (dark grey), CK-p25_1w_ (1-week, light green), CK-p25_2w_ (2 weeks, green), CK-p25_6w_ (6 weeks, dark green). **B**: Heat map showing the pixel-wise standard variation of the bootstrapped persistence images across control and CK-p25 conditions. **C:** Schematic for filtering persistence barcodes with MorphOMICs. Starting from microglial rooted tree, only bars are selected that are born at 0µm and independent of their length (representing likely primary branches). These bars are converted into a persistence diagram. **D-F**: Palantir trajectory of all brain region (without cochlear nucleus and cerebellum for simplicity) from control and CK-p25 condition with highlighted FC microglia trajectory for females (left) and males (right) with (**D**) unfiltered or filtered bars (**E**: start radial distance from the soma = 0µm, **F**: maximum bar length = 10µm). * p < 0.05, ** p < 0.01.

To identify which properties are potentially relevant, we looked at the most variable pixels across CK-p25 FC_mg_ and the control bootstrapped persistence images (**Fig. 7C**). We found the highest variability along the diagonal and close to origin of the persistence diagram corresponding to short branches and branches close to the soma (see also **Fig. 1B**, v). We therefore decided to zoom in on the short and the long persistence bars, filtered them out separately, and repeated our MorphOMICs analysis (**Fig. 7D**). Using this method, we saw that just a few long bars sufficed to capture the sexually dimorphic phenotypes along the disease trajectory (**Fig. 7E, F**). In contrast, we observed a different dimorphic control-to-disease spectrum if we focused only on the short bars (**Fig. 7G**). These results suggest that persistence barcodes highlight different phenomena, and therefore both short and long bars are essential for the understanding of morphology.

## Discussion

In this study, we analyzed heterogeneity and sexual dimorphism of microglia morphology across seven brain regions from a total of 39,190 cells through development and disease (**Supp. Table 1**). To do so, we developed and applied the MorphOMICs pipeline, which extracts the information of the entire reconstructed microglial tree in a minimally biased way, combined with variability reduction and improved data visualization.

### Insights into the microglial phenotypic spectrum

Microglial functional heterogeneity has been indicated using brain-region-specific single-cell transcriptome analysis (Furube et al., 2018; Grabert and McColl, 2018; Mildner et al., 2011), but morphological differences have been difficult to identify and quantify. MorphOMICs revealed that the microglia in an adult brain exhibit regional heterogeneity (**Fig. 1F**) that exists already in early postnatal development (**Fig. 3C**) and diminishes during degeneration (**Fig. 4B**, **5B**). Although microglia display a phenotypic spectrum (**Fig. 6**), they respond to disease in a brain-region-specific manner, dependent on the degeneration model. In fact, frontal brain regions are known to be first responders in Alzheimer-like degeneration (Bakkour et al., 2013; Desikan et al., 2010).

Even though we did not directly compare the functional consequences of environmental changes on microglia, we observed some important links. Microglia shifted from the P7/P22 trajectory across all brain regions at P15 (**Fig. 3E**), which is the time of circuit refinement, where microglia have frequently been shown to participate in the process of synaptic pruning (Hattori and McGeer, 1973; Kroon et al., 2019; Maklad and Fritzsch, 2003; Wong et al., 2005). Another synapse-associated pattern occurred in the DG_mg_ of CK-p25_2w_ (**Fig. 5D**), where we unexpectedly found the highest CD68 fold change and not within the CK-p25_6w_, where we have observed the most distinct morphological shift from the control. This discrepancy might be associated with transient p25 expression, which has been previously described (Fischer et al., 2005). In general, we could associate CD68 upregulation with an increasingly diverse phenotype, indicating an increased reactivity, as exhibited in ♀FC_mg_ CK- p25. However, this effect was less prominent in ♂FC_mg_ (**Fig. 5D**), suggesting that CD68 upregulation is sex-, brain-region-, and context-dependent. This aspect has to be taken into account for future use of CD68 as a reactivity marker.

### Sex as confounding factor of microglial phenotypic spectrum

To what extent sex affects microglial morphology has long been debated (Lenz et al., 2013; Nelson et al., 2017). Here, we confirmed that a sex-specific phenotype exists, which as we have shown is rather mild during adulthood (**Fig. 2B**) but prominent during development (**Fig. 3D**) and degeneration (**Fig. 4C**, **5C**). We found a sexually dimorphic microglial response in both degeneration models, which was pronounced in the immediately affected brain regions FC_mg_, DG_mg_, and S1_mg_. Interestingly, females showed an earlier shift along the morphological spectrum compared to males. This supports studies that have suggested a sex-dependent difference in Alzheimer’s disease progression (Congdon, 2018; Gamache et al., 2020; Manji et al., 2019) and points to females having a higher risk of developing dementia (Payami et al., 1996; Turner, 2001).

Estrogens have been shown to be involved in the masculinization of the brain (Lenz and McCarthy, 2015; Lenz et al., 2013; Nissen, 2017), and microglia are also suspected of playing a role in this process (Lenz and McCarthy, 2015; Lenz et al., 2013). We were surprised to see that microglia from the ovariectomized females were distinct from their non-ovariectomized counterparts, but also that brain regions intermingled differently (**Fig. 2B-C**). Whereas ♀FC_mg_ and ♀OB_mg_ occupied a similar cluster in control adults, ♀_ov_FC_mg_ were distinct from ♀_ov_OB_mg_ and highly intermingled with ♀_ov_SN_mg_ (**Fig. 2C**), suggesting that the impact of estrogens on microglial morphology is complex. Overall, MorphOMICs links the previously reported sexually dimorphic microglial transcriptome in the healthy brain (Ayata et al., 2018; Schwarz et al., 2012; Thion et al., 2018; Villa et al., 2018) and in degeneration models (Bruce-Keller et al., 2000; Crain and Watters, 2010; Vegeto et al., 2006; Villa et al., 2015, 2016; Yanguas-Casás et al., 2018) with a distinct morphological phenotype.

### MorphOMICs as a novel tool for morphological analysis

An advantage of MorphOMICs is that it preserves the intrinsic properties of the reconstructed morphological tree and avoids feature-selection-derived biases. Interestingly, we found that both short and long bars within a barcode contain information that contribute to microglial spectrum (**Fig. 7E-G**). Future studies will focus on identifying particularly informative regions of a persistence barcode, which provides a perspective for morphological analysis of lower-resolution images, such as *in vivo* microglial imaging for potential non-invasive diagnostic applications. Stable ranks would provide a mathematically robust approach to address this question, as we have shown already that standard stable ranks of the TMD captured the microglial phenotypes as well as the persistence images of the microglial TMD (**Supp. Fig. 2G**) (Riihimäki and Chacholski, 2018).

MorphOMICs provides an advanced strategy for systematically comparing microglial populations across different brain regions and conditions: this could be expanded infinitely. We therefore provide our dataset in NeuroMorpho.org and the code as a download on GitHub (https://git.ist.ac.at/rcubero/morphomics), with detailed instructions on implementation. A critical point to consider is the number of cells that are needed for MorphOMICs. While we identified a suitable bootstrap size in **Supp. Fig. 2D**, the condition-specific variability in microglial morphology needs to be systematically assessed to determine the minimum cell number before MorphOMICs can be reliably applied.

Most important, MorphOMICs overcomes the dichotomized view of microglial morphology to either ramified, relating to a surveilling function, or amoeboid, for highly phagocytosing. This is relevant because microglia present a wide spectrum of morphological responses to local cues that are neglected with classical analysis, thus compromising the use of morphology as a readout of environmental modification.

## Supplementary Figures

**Supplementary Figure 1.**
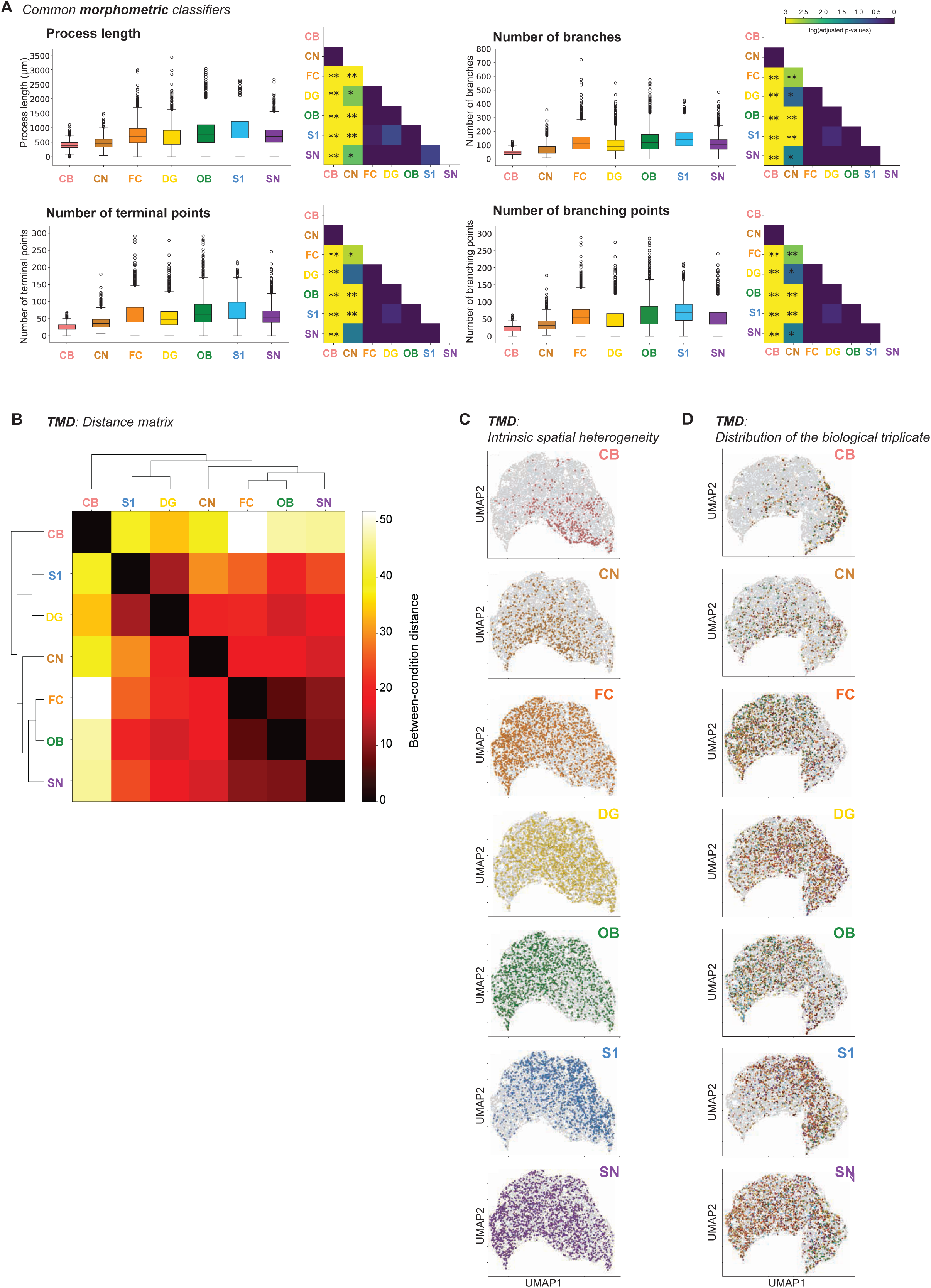
Classic morphometry analysis and intrinsic variability of microglial morphology. **A:** Box plots for the following morphometric features: process length, number of branches, and terminal and branching points for traced adult C57BL/6J microglia across different brain regions: CB (cerebellum, n=299), CN (cochlear nucleus, n=498), FC (frontal cortex, n=926), DG (dentate gyrus, n=902), OB (olfactory bulb, n=796), S1 (somatosensory cortex, n=719), SN (substantia nigra, n=1050) from at least six animals. Next, matrices with color-coded p-values for the pairwise comparison of each morphometric (see also **Supp. Table 2**). **B:** Hierarchically-ordered heat map for pairwise TMD intrinsic distances between average persistence images from microglia across brain regions from Fig. 1A. **C**-**D:** UMAP plots of the entire microglial population size (grey) with color-highlighted brain regions (**C**) or animals (**D**). Each dot represents a single persistence image. (**D**) Triangle and circle for females and males, respectively. * p < 0.05, ** p < 0.01.

**Supplementary Figure 2.**
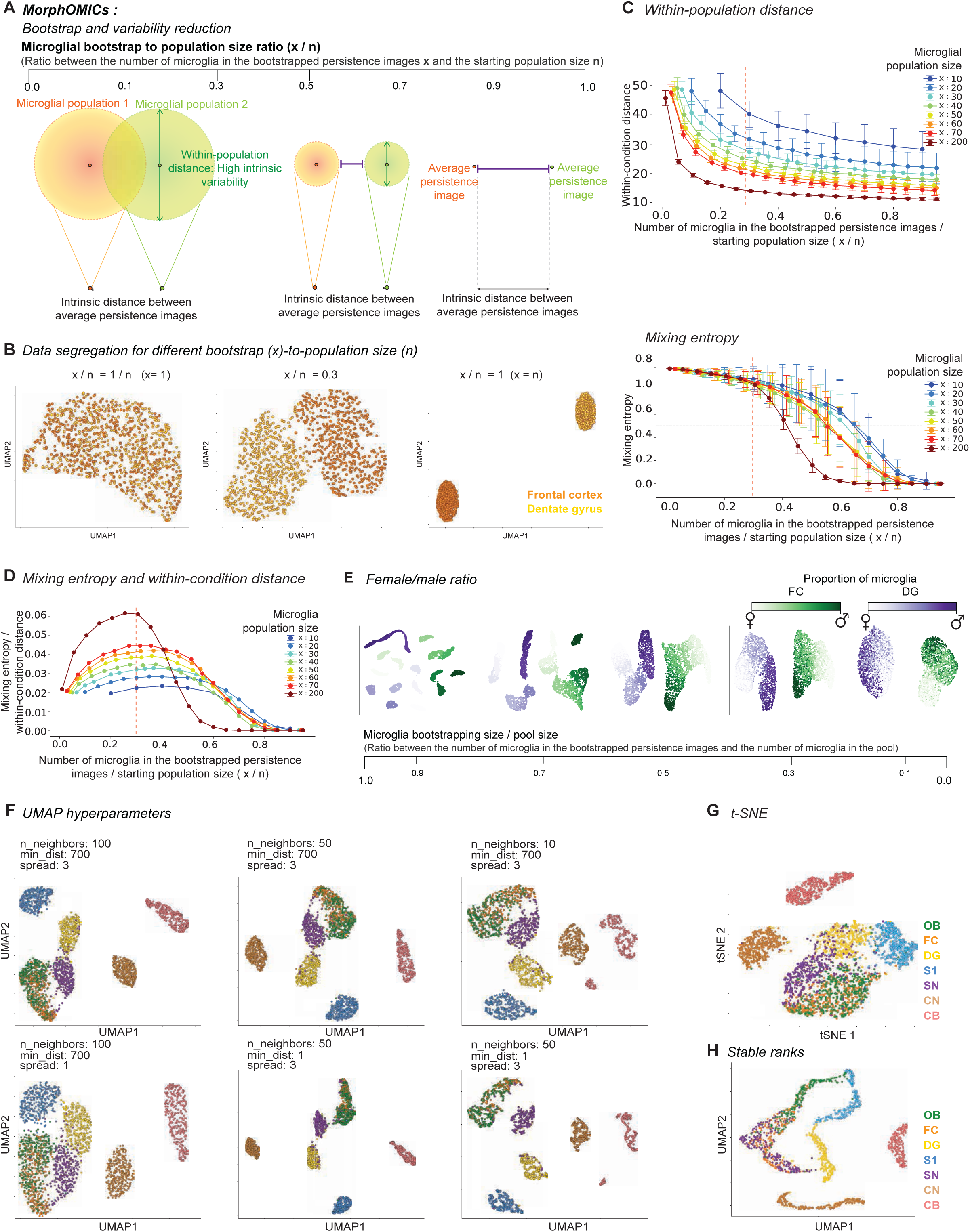
Details about the MorphOMICs paradigm. **A:** Schematic of the bootstrapping effects on the distance between tree structures from the same population (within-population distance, green arrows) and two distinct populations (distance between average persistence images, purple lines). Increase of bootstrap-to-population size ratio (x/n) reduces within-population distance and increases distance between average persistence images. **B:** UMAP plots of MorphOMICs-analyzed microglia for frontal cortex (orange) and dentate gyrus (yellow) for different bootstrap-to-population size ratios. Left: x=1, allows no segregation. Middle: x/n=0.3. Right: x/n=1 causes accentuation. **C:** Line plot ± SD displays how within-population distance (top) and mixing entropy (bottom) decrease with enhanced bootstrap-to-population size ratio (x/n). An empirical threshold of 0.3 was selected (red dashed line). **D:** Line plot ± SD displays how the ratio between mixing entropy and within-condition distance varies by enhancing the bootstrap-to-population size ratio (x/n). **E:** UMAP plots of MorphOMICs-analyzed microglia for frontal cortex (green) and dentate gyrus (purple) for different bootstrap-to-population size ratios and varying male-to-female ratios within the population size. Olfactory bulb (OB), frontal cortex (FC), dentate gyrus (DG), somatosensory cortex (S1), substantia nigra (SN), cochlear nucleus (CN), and cerebellum (CB). **F:** UMAP plots of MorphOMICs-analyzed microglial morphology across seven brain regions as shown for Fig. 1F with varying hyperparameters for number of neighbors (n_neighbors), minimum distance (min_dist), and spread. **G:** t-SNE plot of MorphOMICs-analyzed microglia across seven brain regions as shown in Fig. 1F for UMAP. **H:** UMAP plots of stable-rank representation of microglial morphology (see Methods: *Stable Ranks*) across seven brain regions. SD: standard deviation.

**Supplementary Figure 3.**
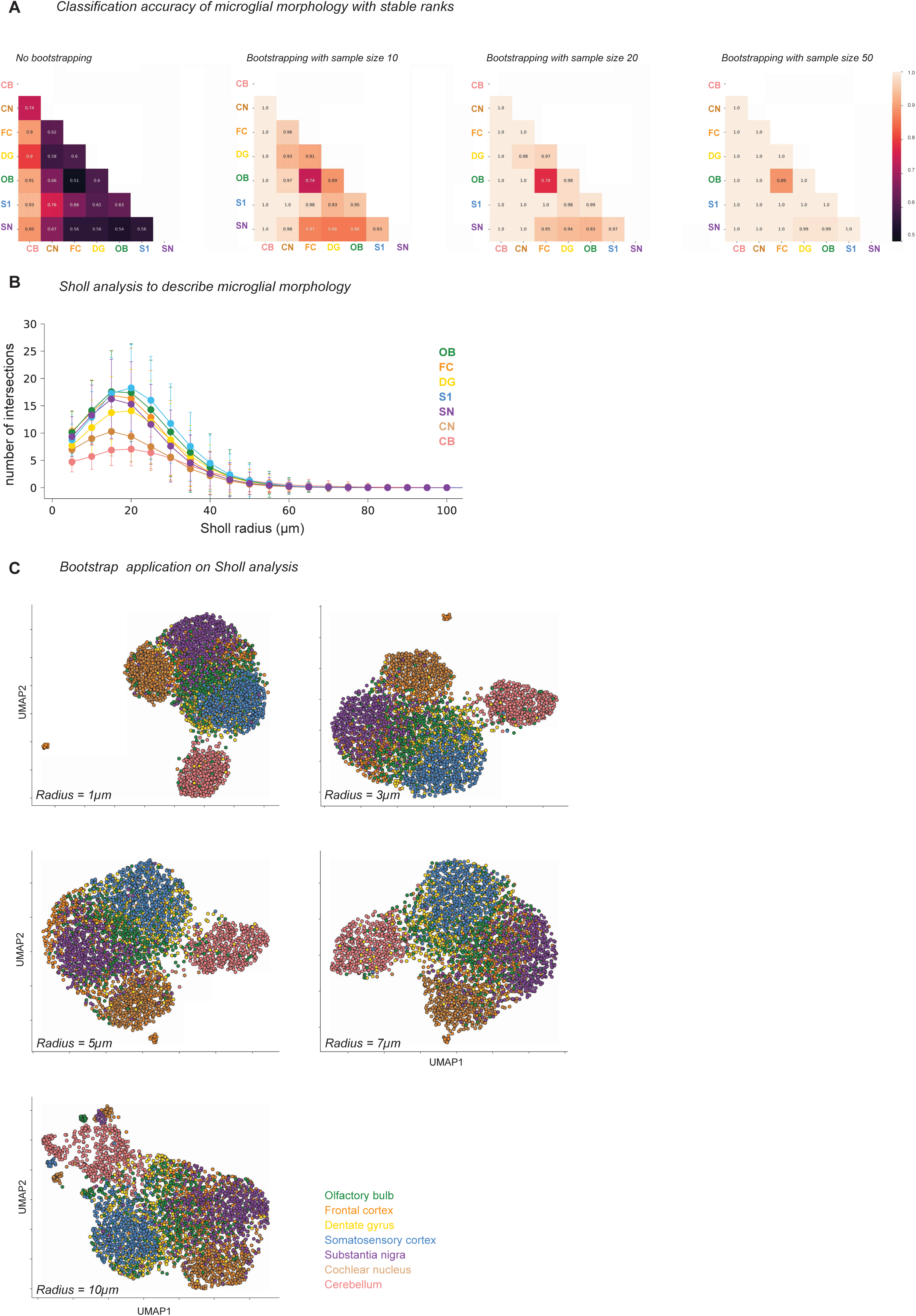
Sholl analysis of microglial heterogeneity depends on the Sholl step size radius. **A:** Heatmap of classification accuracy between pairs of brain regions using stable ranks for different bootstrap sizes. Numbers indicates the percentage of microglia correctly assigned in the classification task, averaged over 10 repeated cross-validations. 1, perfect assignment; 0.5 random assignment. **B:** Sholl curves showing the number of processes, averaged per each control brain region ± SD, that intersect with a series of concentric Sholl spheres spaced at 5µm. **C:** UMAP plot of Sholl-analyzed microglia, color-coded for each brain region. Each dot represents a bootstrapped Sholl analysis, setting the consecutive radius step size at 1, 3, 5, 7, 10 µm respectively. SD: standard deviation.

**Supplementary Figure 4.**
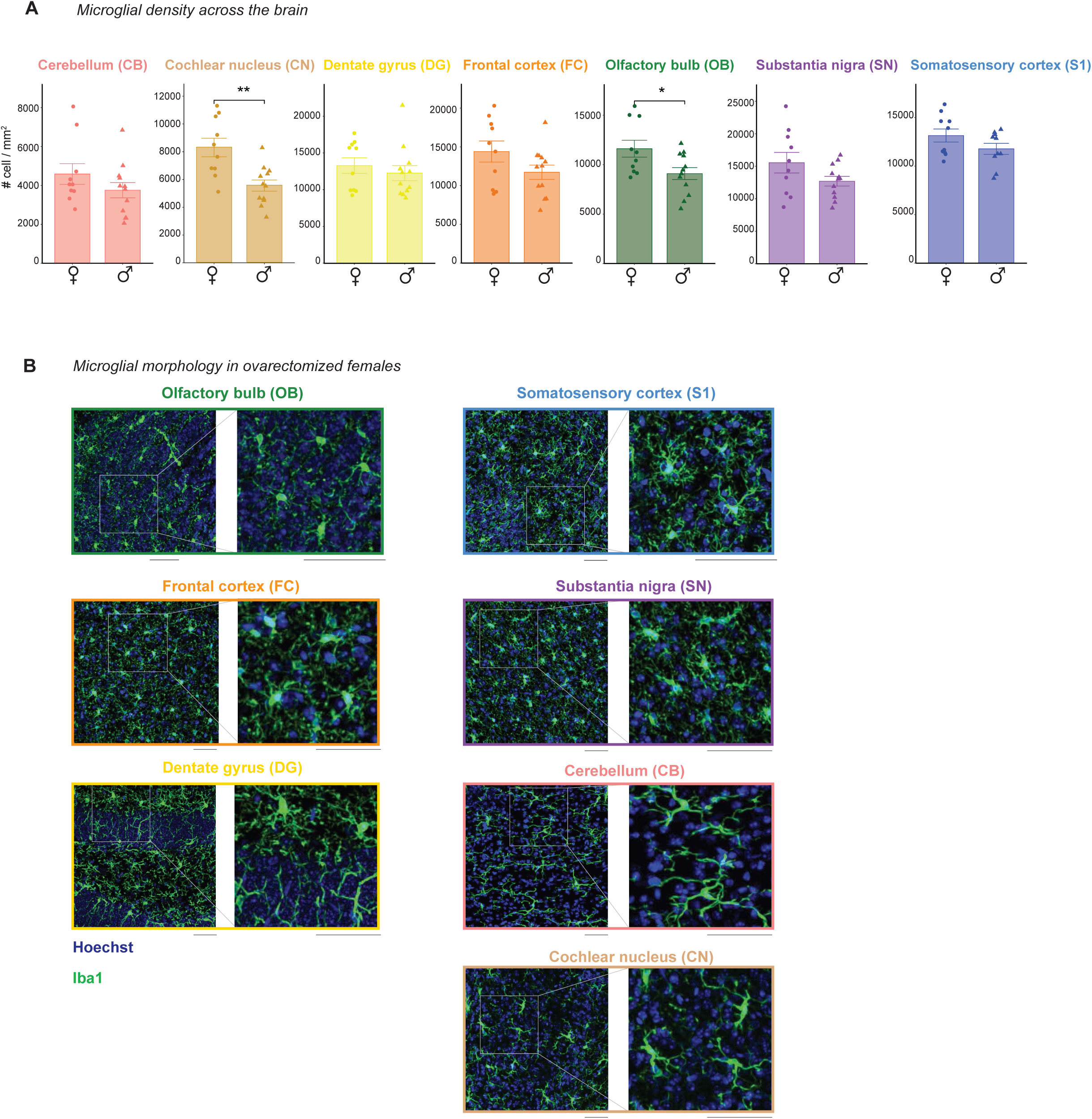
Adult microglial density and the impact of ovariectomy on their phenotype. **A:** Bar plot describing the distribution of microglial density across brain regions in C57BL/6J female and male adults. Cell density is expressed as the number of cells per mm^2^ ± SD. Sex averages for microglia from each region were compared with two-sided t-test. CN_mg_ (♀=11, ♂= 11; t = 3.504, df = 15.1, p-value = 0.00317) and OB_mg_ (♀=11, ♂= 11; t = 2.401, df = 16.864, p-value = 0.0282). **B:** Confocal images of immunostained microglia (Iba1, green) and cell nuclei (Hoechst, blue) from ovariectomized C57BL/6J adult mice for each brain region with zoom-in. Scale bar: 50 μm.

**Supplementary Figure 5.**
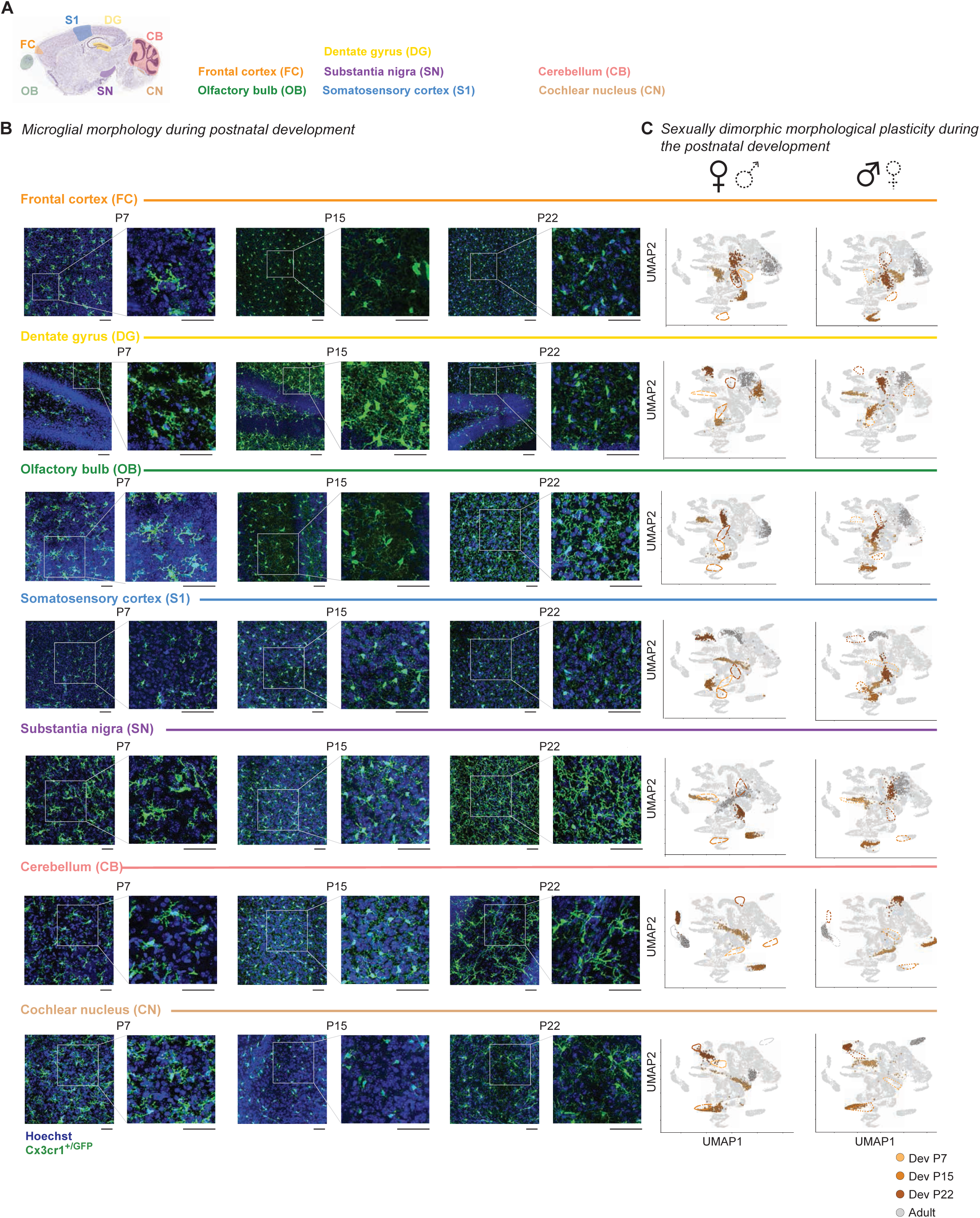
Microglial phenotypic spectrum during postnatal development. **A:** Sagittal view of analyzed brain regions: olfactory bulb (OB), frontal cortex (FC), dentate gyrus (DG), somatosensory cortex (S1), substantia nigra (SN), cochlear nucleus (CN) and cerebellum (CB). (Image credit: Allen Institute (Lein et al., 2006)) **B:** Confocal images of GFP^+^ (green) microglia and cell nuclei (Hoechst, blue) from Cx3cr1^+/GFP^ mice at P7, P15, and P22 for each brain region with zoom-in. Scale bar: 50 μm. **C:** UMAP plots of MorphOMICs-analyzed microglia from **B** for females (left) and males (right) at P7 (light brown), P15 (orange), P22 (dark brown), and adults (dark grey). Separate UMAP for each brain region and sex. Each dot represents a bootstrapped persistence image.

**Supplementary Figure 6.**
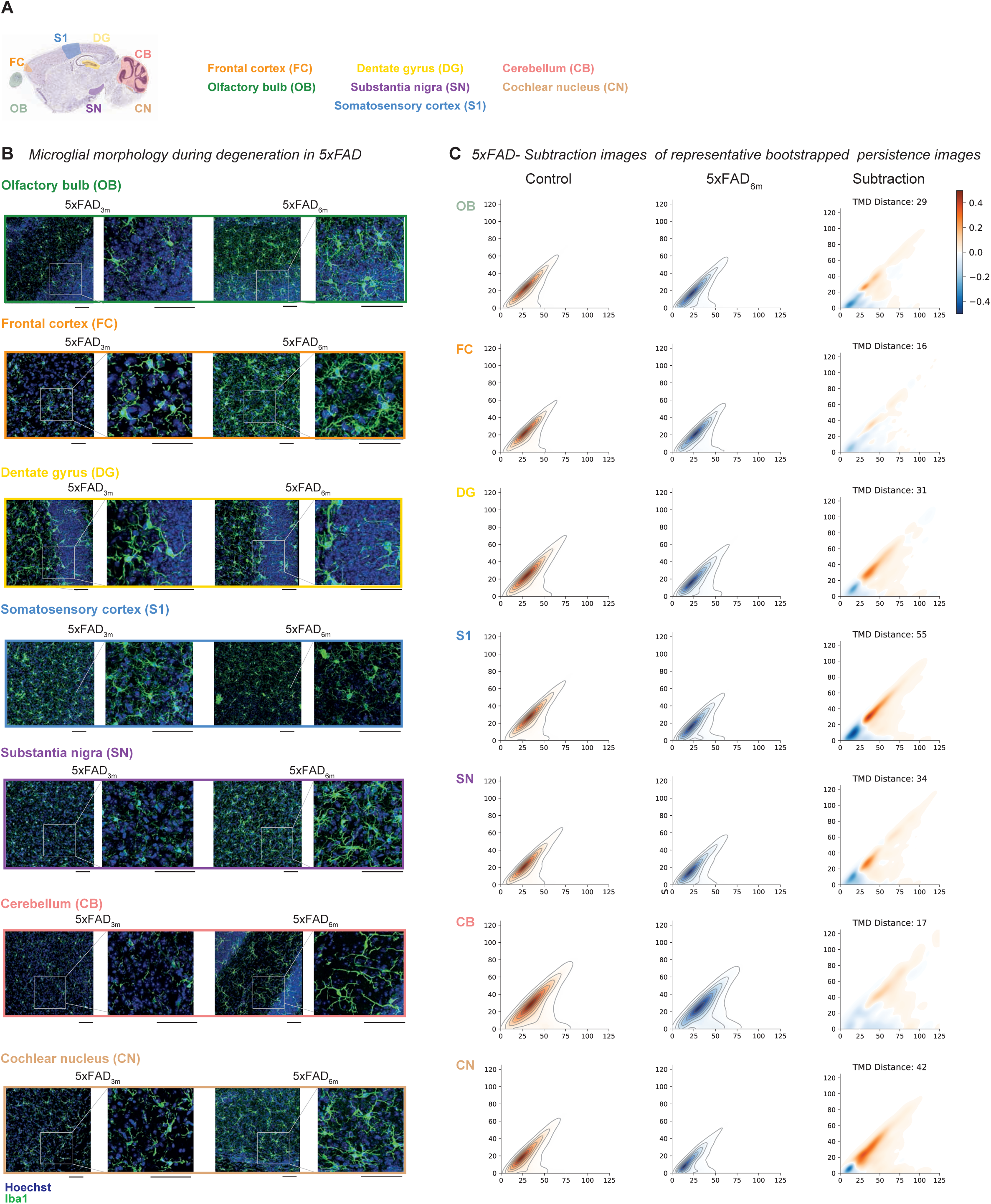
Microglia phenotypic spectrum in the 5xFAD model of familiar Alzheimer’s neurodegeneration. **A:** Sagittal view of analyzed brain regions: olfactory bulb (OB), frontal cortex (FC), dentate gyrus (DG), somatosensory cortex (S1), substantia nigra (SN), cochlear nucleus (CN) and cerebellum (CB). (Image credit: Allen Institute (Lein et al., 2006)) **B:** Confocal images showing stained microglia (Iba1, green) and cell nuclei (Hoechst, blue) from the analyzed brain regions in 5xFAD_3m_ and 5xFAD_6m_ (3 and 6 months, respectively) with zoom-in. Scale bar: 50 μm. **C:** Representative persistence images corresponding to each cluster centroid from Fig. 4B with color-coded process density (orange = control; blue = 5xFAD_6m_) and respective subtraction images per every brain region (orange = increased control process density; blue = increased 5xFAD_6m_ process density)

**Supplementary Figure 7.**
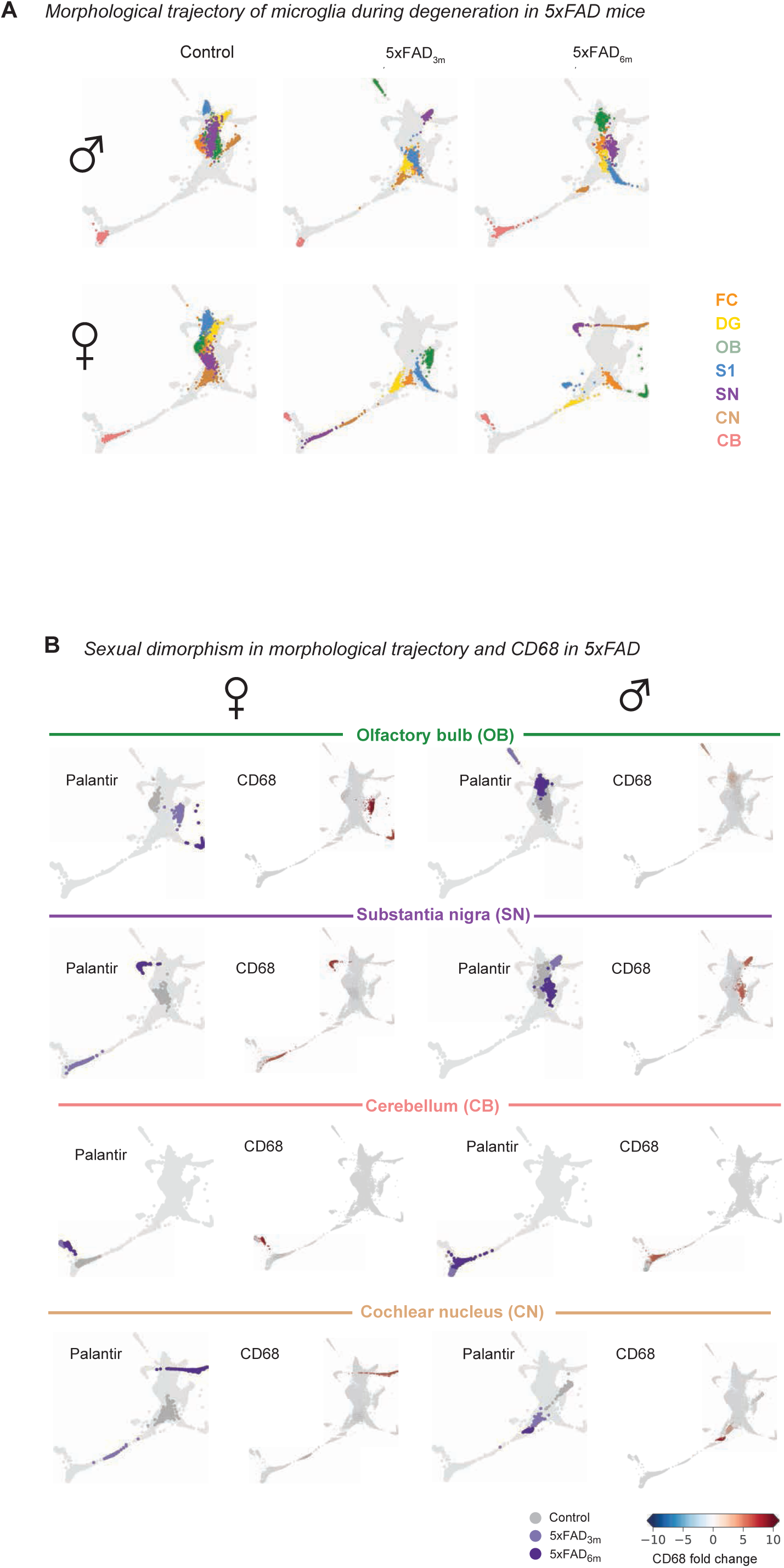
Sexually dimorphic microglia phenotype in 5xFAD. **A:** Palantir reconstruction of microglial morphological trajectory in males (top) and females (bottom) from **Supp. Fig. 6B**. Each time-point highlighted in a separate Palantir plot. **B:** Palantir reconstruction of microglial trajectory with corresponding color-coded CD68 fold change next to it for females (left) and males (right) in control adult, 5xFAD_3m_, and 5xFAD_6m_ of OB, SN, CN, and CB. Fold change < 0 blue; > 0 red.

**Supplementary Figure 8.**
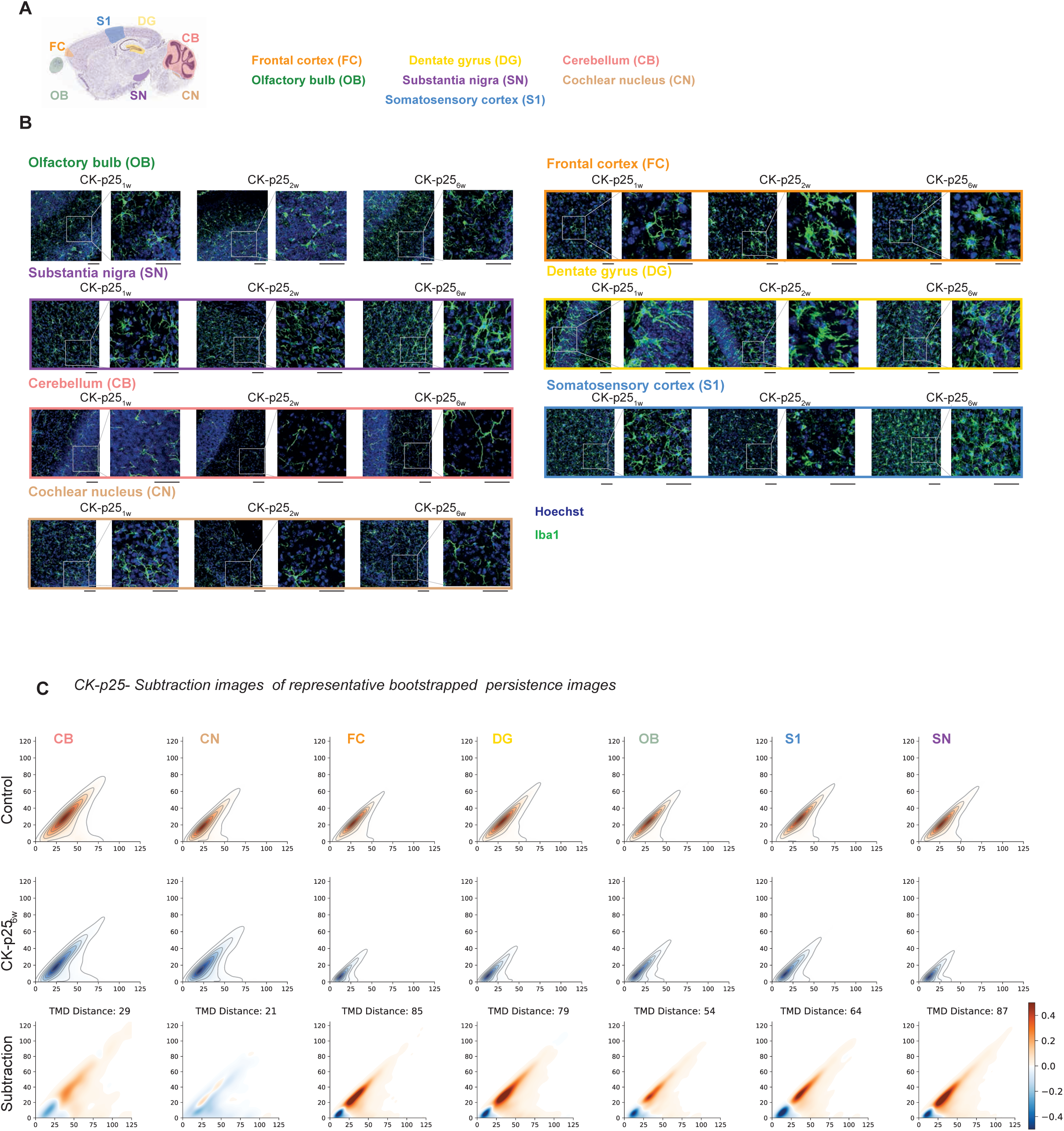
Microglia phenotypic spectrum in the CK-p25 model of sporadic neurodegeneration. **A:** Sagittal view of analyzed brain regions: olfactory bulb (OB), frontal cortex (FC), dentate gyrus (DG), somatosensory cortex (S1), substantia nigra (SN), cochlear nucleus (CN) and cerebellum (CB). (Image credit: Allen Institute (Lein et al., 2006)) **B:** Confocal images showing stained microglia (Iba1, green) and cell nuclei (Hoechst, blue) from analyzed brain regions and CK-p25_1w_, CK-p25_2w_, CK-p25_6w_ mice (1-, 2- and 6-weeks after doxycycline withdrawal) with zoom-in. Scale bar: 50 μm. **C:** Representative persistence images corresponding to each cluster centroid from Fig. 5B with color-coded process density (orange = control; blue = CK-p25_6w_) and respective subtraction images per every brain region (orange = increased control process density; blue = increased CK-p25_6w_ process density)

**Supplementary Figure 9.**
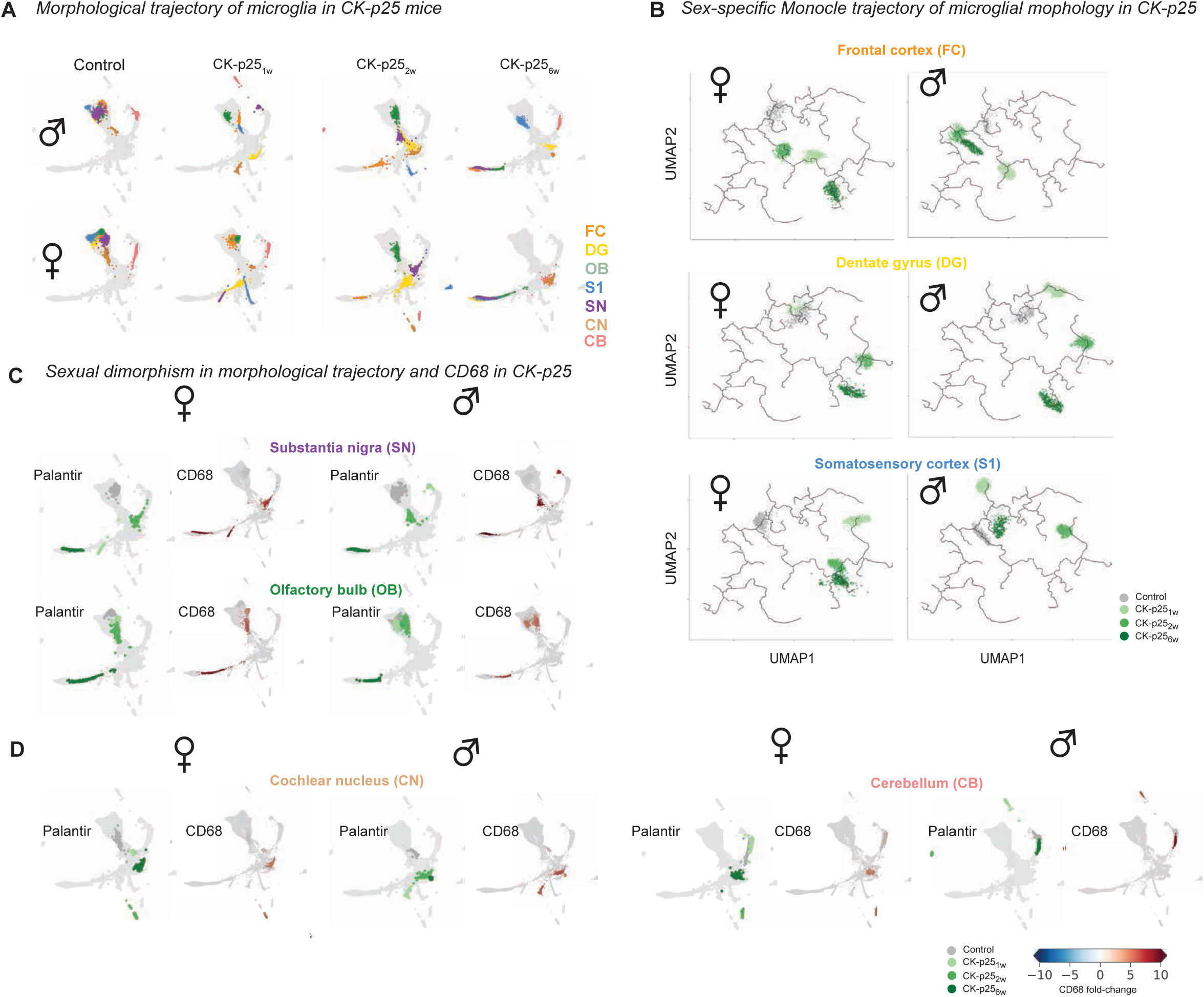
Sexually dimorphic microglial phenotype in CK-p25. **A:** Monocle reconstruction of microglia trajectory of females (left) and males (right) in control, adult and CK-p25 mice FC, DG, and S1. **B:** Palantir reconstruction of microglial morphological trajectory in males (top) and females (bottom) in control and CK-p25 mice. Each disease time-point highlighted in a separate Palantir plot. **C**-**D:** Palantir reconstruction of microglia trajectory and corresponding color-coded CD68 fold-change of females (left) and males (right) from **Supp. Fig. 8B** for SN, OB (**C**) and CN, CB (**D**). Fold change < 0 blue; > 0 red.

**Supplementary figure 10.**
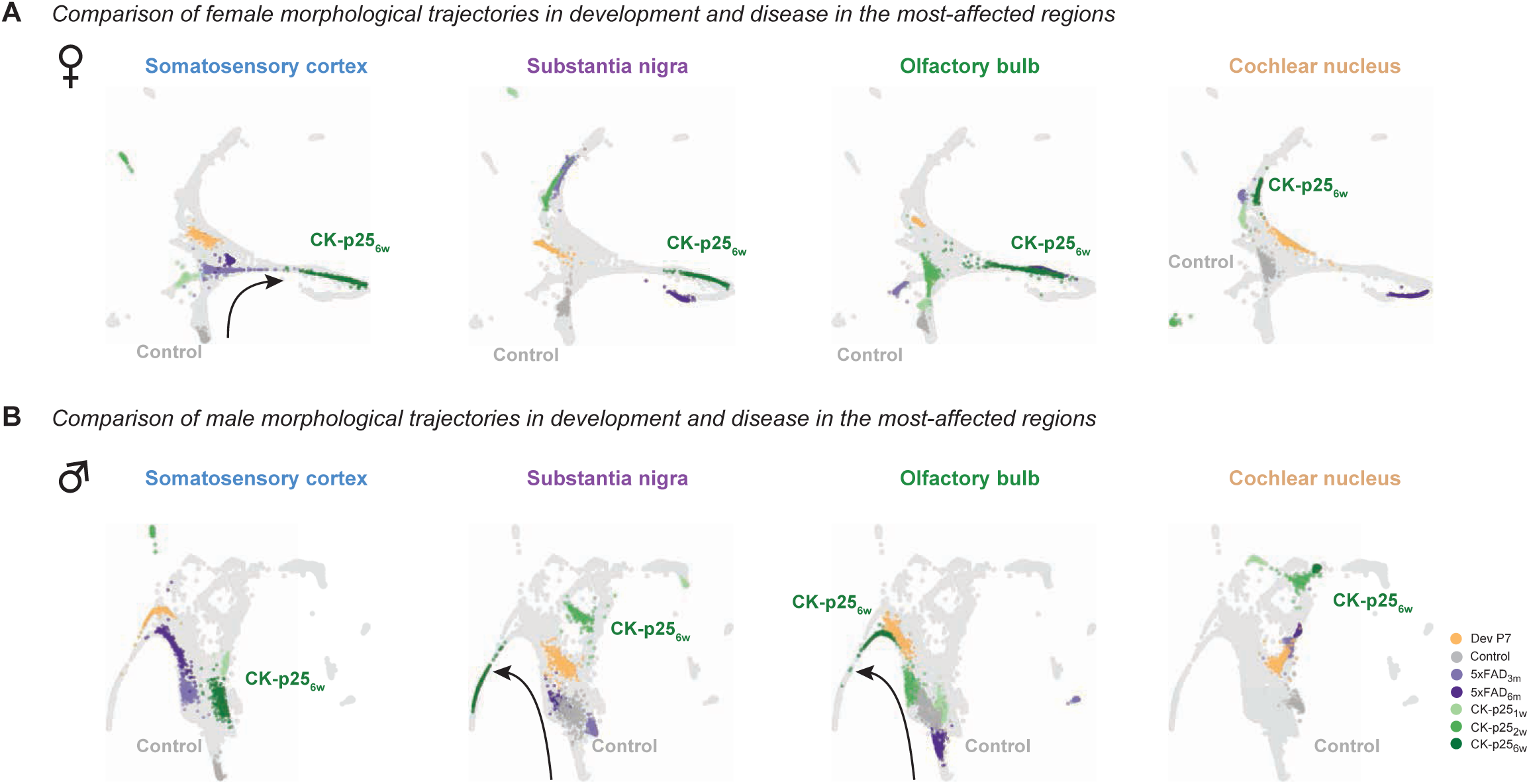
Integration of early developmental phenotype into the disease spectrum in secondarily-affected regions. **A**-**B:** Palantir reconstructions of microglia trajectory in females (**A**) and males (**B**) for S1, SN, OB, and CN in control (dark grey), P7 (light brown), CK-p25_1w_ (1-week, light green), CK-p25_2w_ (2 weeks, green), CK-p25_6w_ (6 weeks, dark green), 5xFAD_3m_ (3 months, light purple), and 5xFAD_6m_ (6 months, purple) conditions. Each brain region is highlighted in a separate Palantir plot. Black arrow: control-to-disease spectrum

**Supplementary figure 11.**
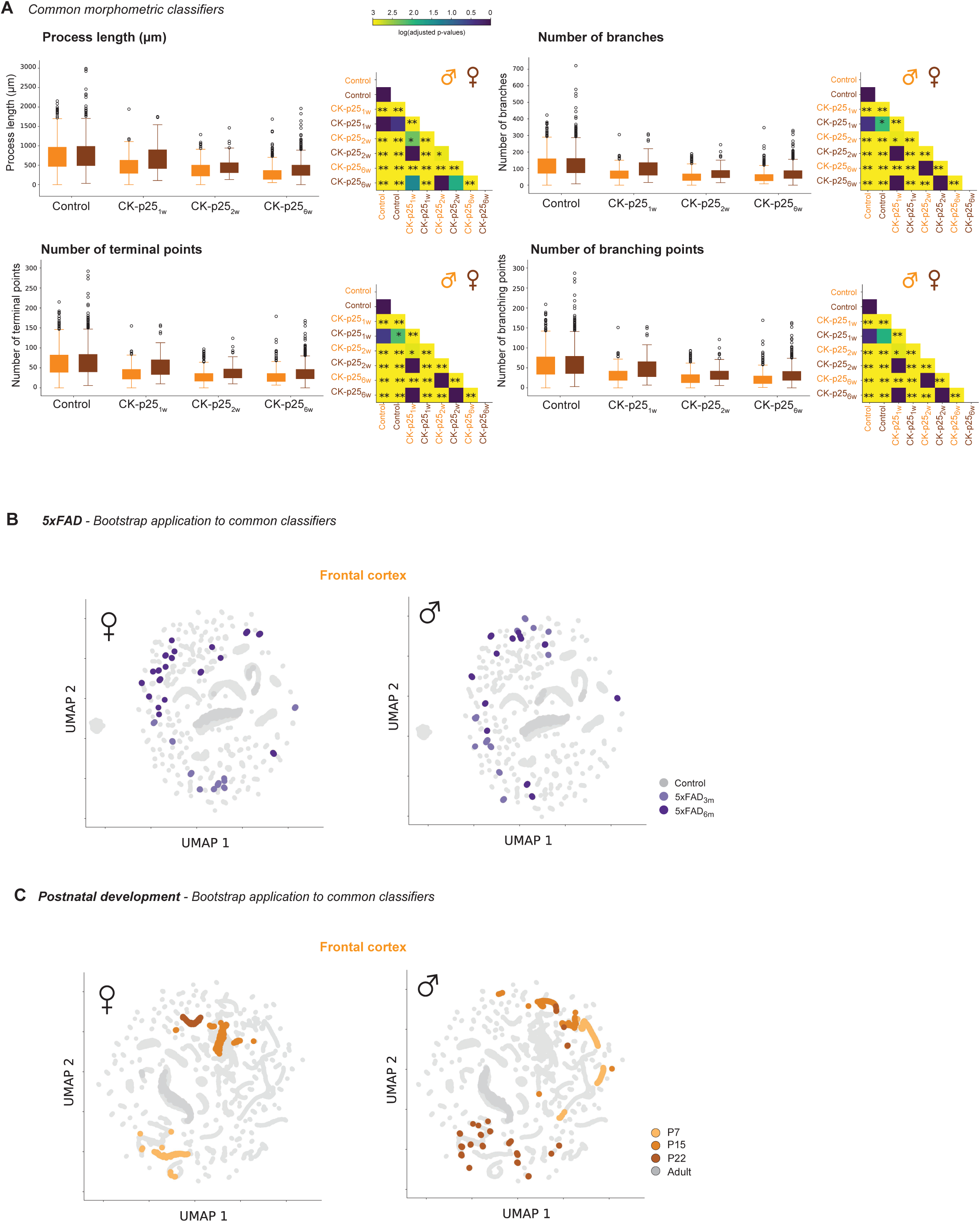
Classical morphometric do not recapitulate MorphOMICs observations. **A:** Left: box plots for the selected features dendritic length, number of branches, and terminal and branching points of control (n_♀_= 926, n_♂_= 894), and CK-p25_1w_ (n_♀_= 219, n_♂_= 194), CK- p25_2w_ (n_♀_= 264, n_♂_= 492), CK-p25_6w_ (n_♀_= 858, n_♂_= 462) mice (1-, 2- and 6-weeks after doxycycline withdrawal, respectively) in the frontal cortex (FC). Right: matrices showing color-coded p-values for the pairwise comparison of each morphometric. **B**-**C:** UMAP representations based on an extended list of morphometrics (see **Supplementary Table 4.**) applied to females (left) and males (right) in 5xFAD (**B**) and Cx3cr1-GFP^+/-^ mice (**C**). Highlighted clusters are FC for control/adult (dark grey), 5xFAD_3m_ (3 months, light purple), 5xFAD_3m_ (6 months, dark purple), P7 (light brown), P15 (orange) and P22 (dark brown) respectively.

**Supplementary Table 1.**
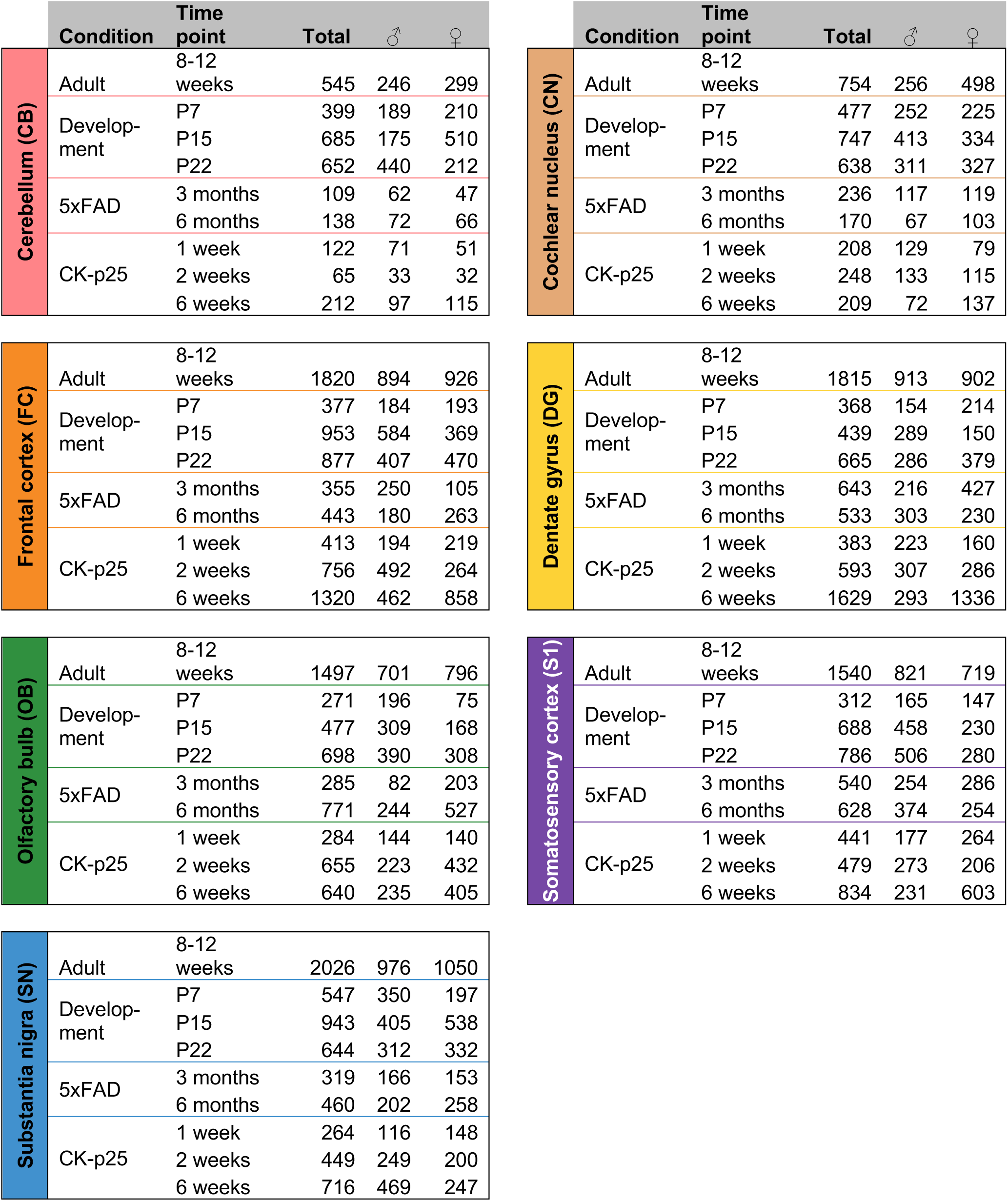
Number of traced microglia for each brain region, sex, and condition.

**Supplementary Table 2.**
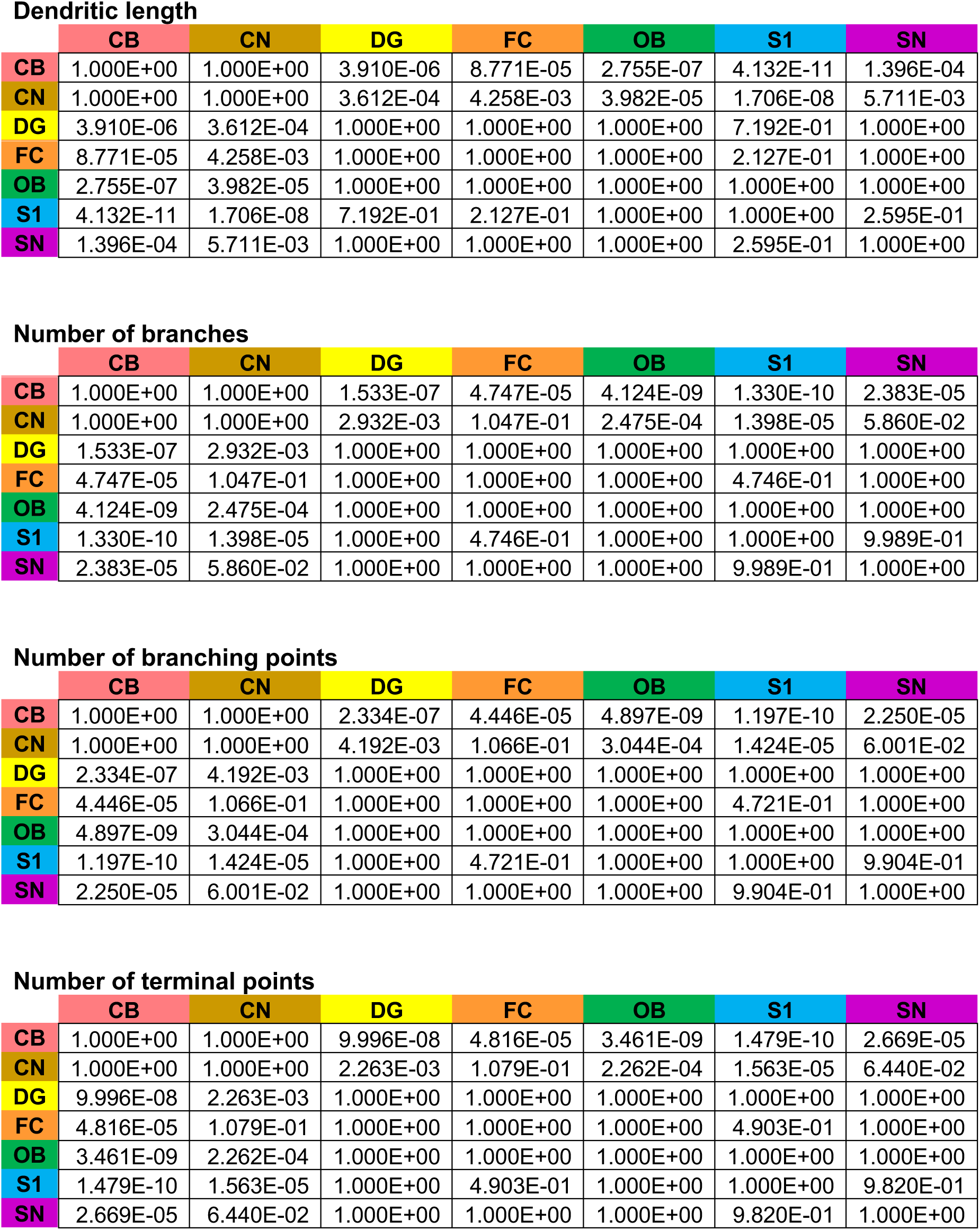
Statistical tests related to Supplementary Figure 1A. P-value for pairwise comparisons between adult microglia of a brain regions for each morphometric. Cerebellum (CB), cochlear nucleus (CN), dentate gyrus (DG), frontal cortex (FC), olfactory bulb (OB), somatosensory cortex (S1), substantia nigra (SN).

**Supplementary Table 3.**
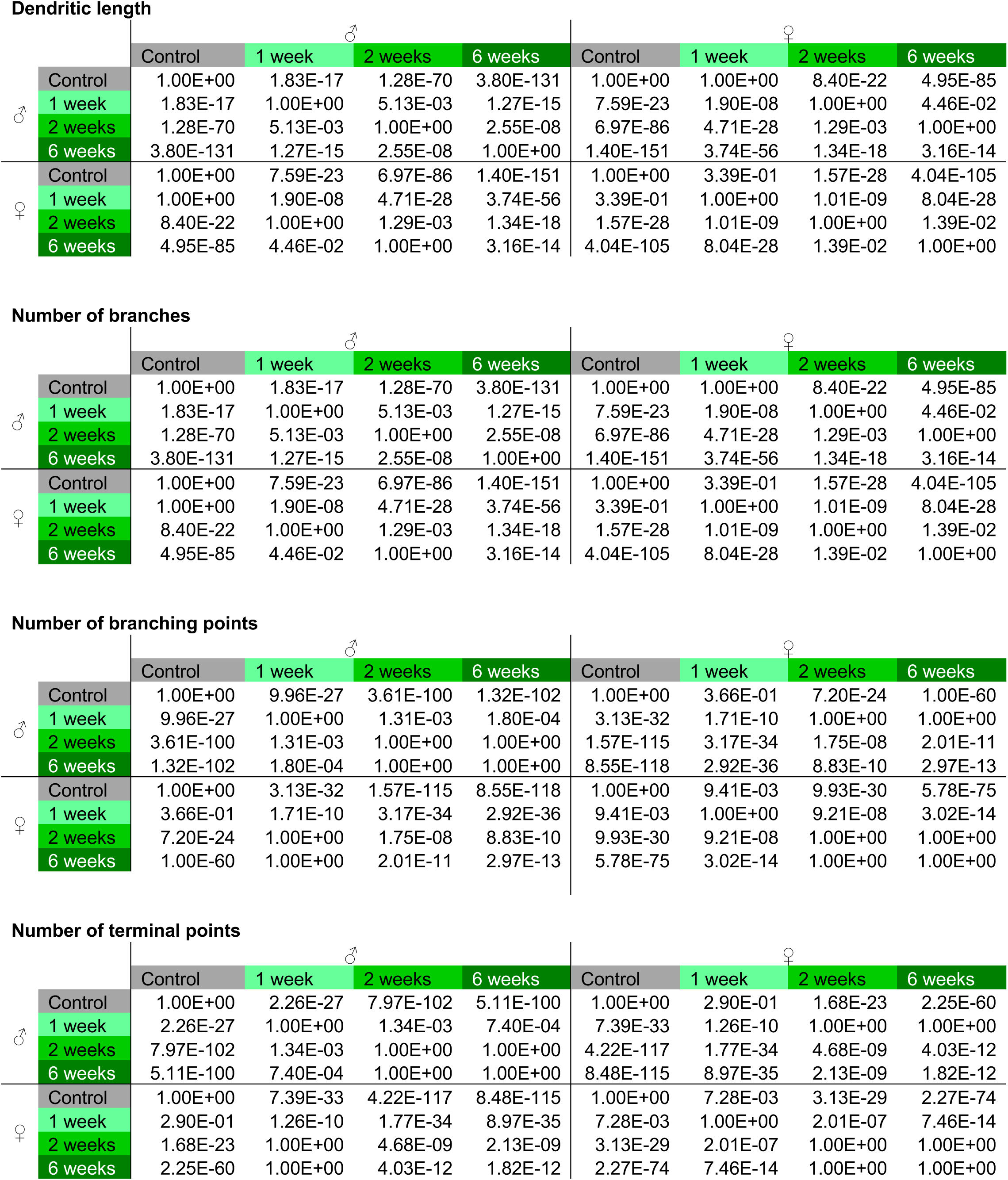
Statistical tests related to Figure 7A. P-value for pairwise comparison between microglia from the frontal cortex of the CK-p25 for different conditions and sex.

**Supplementary Table 4.**
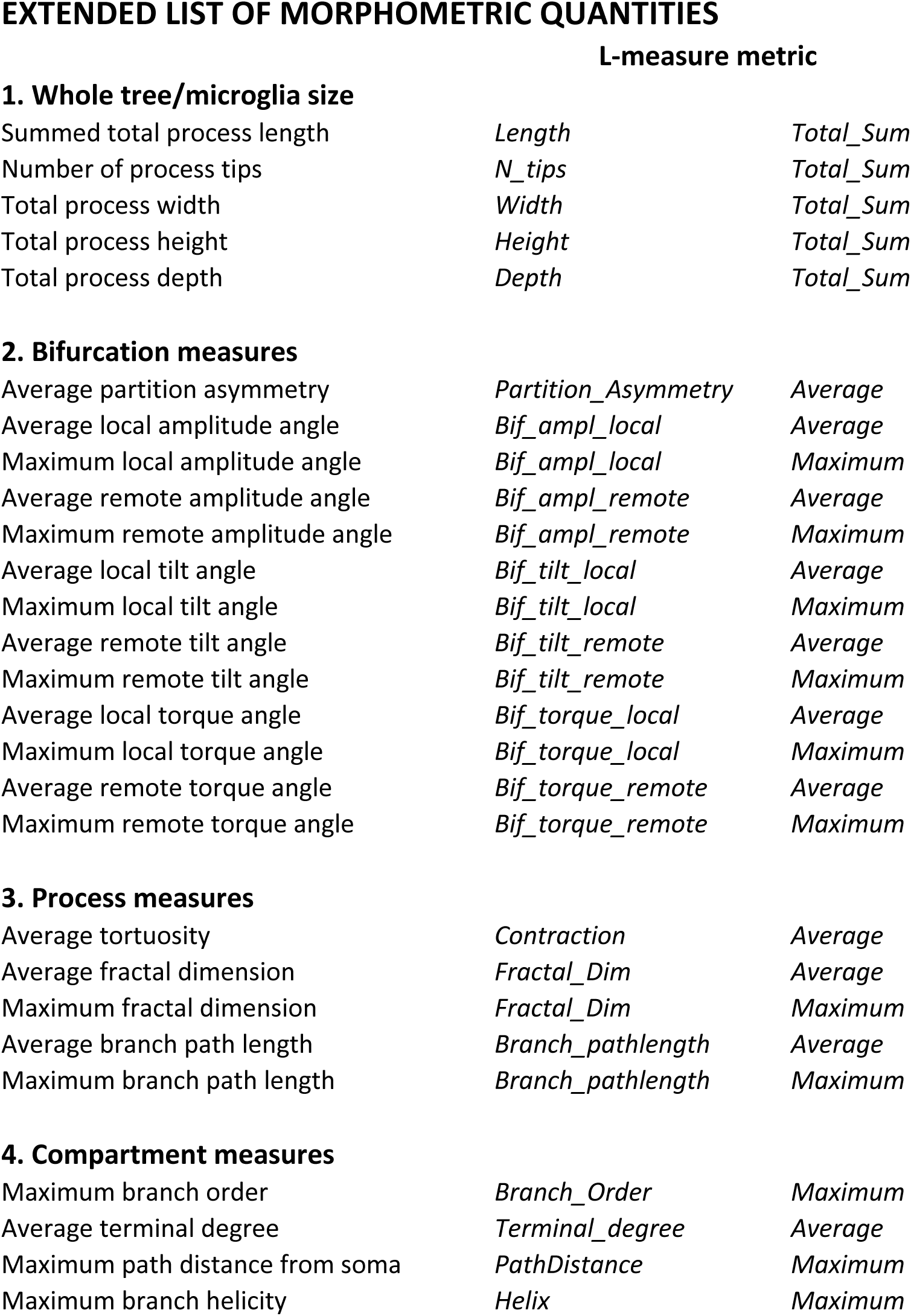
Classical morphometric related to Figure 7A, Supplementary Figure 11B-C. Extended list of classical morphometric quantities (Bijari et al., 2021; Polavaram et al., 2014).

**1. Whole tree/microglia size**
|  |  |  |
| --- | --- | --- |
| Summed total process length | *Length* | *Total_Sum* |
| Number of process tips | *N_tips* | *Total_Sum* |
| Total process width | *Width* | *Total_Sum* |
| Total process height | *Height* | *Total_Sum* |
| Total process depth | *Depth* | *Total_Sum* |
| **2. Bifurcation measures** |  |  |
| Average partition asymmetry | *Partition_Asymmetry* | *Average* |
| Average local amplitude angle | *Bif_ampl_local* | *Average* |
| Maximum local amplitude angle | *Bif_ampl_local* | *Maximum* |
| Average remote amplitude angle | *Bif_ampl_remote* | *Average* |
| Maximum remote amplitude angle | *Bif_ampl_remote* | *Maximum* |
| Average local tilt angle | *Bif_tilt_local* | *Average* |
| Maximum local tilt angle | *Bif_tilt_local* | *Maximum* |
| Average remote tilt angle | *Bif_tilt_remote* | *Average* |
| Maximum remote tilt angle | *Bif_tilt_remote* | *Maximum* |
| Average local torque angle | *Bif_torque_local* | *Average* |
| Maximum local torque angle | *Bif_torque_local* | *Maximum* |
| Average remote torque angle | *Bif_torque_remote* | *Average* |
| Maximum remote torque angle | *Bif_torque_remote* | *Maximum* |
| **3. Process measures** |  |  |
| Average tortuosity | *Contraction* | *Average* |
| Average fractal dimension | *Fractal_Dim* | *Average* |
| Maximum fractal dimension | *Fractal_Dim* | *Maximum* |
| Average branch path length | *Branch_pathlength* | *Average* |
| Maximum branch path length | *Branch_pathlength* | *Maximum* |
| **4. Compartment measures** |  |  |
| Maximum branch order | *Branch_Order* | *Maximum* |
| Average terminal degree | *Terminal_degree* | *Average* |
| Maximum path distance from soma | *PathDistance* | *Maximum* |
| Maximum branch helicity | *Helix* | *Maximum* |

## Material and Methods

### Animals

C57BL/6J (Cat#000664) and B6.129P-Cx3cr1tm1Litt/J (Cat#005582, named here Cx3cr1^GFP/-^, only heterozygous were used) were purchased from The Jackson Laboratories. All animals were housed in the IST Austria Preclinical Facility, with 12 hours light-dark cycle, food and water provided ad libitum. Animal from both sexes were used, as indicated in the results. All animal procedures are approved by the Bundesministerium für Wissenschaft, Forschung und Wirtschaft (bmwfw) Tierversuchsgesetz 2012, BGBI. I Nr. 114/2012, idF BGBI. I Nr. 31/2018 under the numbers 66.018/0005-WF/V/3b/2016, 66.018/0010-WF/V/3b/2017, 66.018/0025-WF/V/3b/2017, 66.018/0001_V/3b/2019, 2020-0.272.234. 5xFAD and CK-p25 mice were obtained from the Tsai lab at MIT. All animal work was approved by the Committee for Animal Care of the Division of Comparative Medicine at the Massachusetts Institute of Technology. 5xFAD mice (B6SJL-Tg(APPSwFlLon,PSEN1*M146L*L286V)6799Vas/Mmjax, Stock No: 34840-JAX) were obtained from Jackson laboratory. CK-p25 mice(Cruz et al., 2003),(Cruz et al., 2006), (Fischer et al., 2005) were generated by breeding CaMKIIα promoter-tTA mice (CK controls) (B6;CBA-Tg(Camk2a-tTA)1Mmay/J, Jackson Laboratory, Stock No: 003010) with tetO-CDK5R1/GFP mice (C57BL/6-Tg(tetO-CDK5R1/GFP)337Lht/J, Jackson Laboratory, Stock No: 005706). CK-p25 mice and their CK control littermates were conceived and raised in the presence of doxycycline-containing food to repress p25 transgene expression. To induce p25 transgene expression, mice were fed a normal rodent diet. p25 transgene expression was induced in adult mice at the age of 3 months. Since it is known that doxycycline acts as inhibitor of microglial activation(Jantzie et al., 2005; Santa-Cecília et al., 2016) and keep microglia in a partially immature phenotype(Erny et al., 2015), we decided to compare brains upon withdrawal of the drug with brains which were never exposed to doxycycline as a control, in order to avoid the overestimation of microglia response. Mice were housed in groups of three to five on a standard 12 h light/12 h dark cycle, and all experiments were performed during the light cycle. Food and water were provided ad libitum.

### Brain samples and analyzed brain regions

We analyzed brains of both sexes from C57BL/6J adult mice (8-12 weeks); Cx3cr1^GFP/-^ mice at postnatal time points P7, P15, P21; 5xFAD mice after 3 and 6 months; and CK-p25 mice 1, 2, and 6 weeks after doxycycline withdrawal. We focused on the following brain regions: the glomerular layer of the olfactory bulb (OB), cortical layer III-V of the frontal cortex (FC) and the primary somatosensory cortex (S1), the dentate gyrus of the hippocampus (DG), the substantia nigra (SN), the cochlear nucleus (CN), and the third lobe of the cerebellum (CB). The sagittal view of the brain sections analyzed (**Fig. 1A, 2A, 3A, 4A, 5A** and **Supp. Fig. 4A and 5A)** was taken from the Allen Developing Mouse Brain Atlas-Sagittal atlas and modified to show brain regions-of-interest (Lein et al., 2006).

### Ovariectomy

Adolescent C57BL/6J females at P20 were anesthetized with 5% isoflurane in 0.5 l/min O_2_ during the anesthesia induction and 2% isoflurane in 0.5 l/min O_2_ during the maintenance phase. Using an electric razor, the fur was shaved to expose the skin over the lumbar spine and the region was sterilized with 70% (v/v) ethanol. A midline incision of approximately 1 cm was made on the skin in the lower back, below the chest. The subcutaneous tissue was gently dissected to expose the muscular fascia, and the ovarian fat-pad was identified under the muscular layer. The peritoneal cavity was cut with a 0.5 cm incision. The Fallopian tube was exposed, and the ovary identified and cut at the level of the oviduct. The blood vessels were cauterized to prevent bleeding. The remaining part of the Fallopian tube was placed back in the peritoneal cavity, and the muscular fascia was sutured. The same protocol was repeated for the contralateral ovary. At the end, the skin was sutured. The animals received Metamizol (Sanofi Aventis, Cat#Ay005, *s.c.* 200 mg/kg during surgery) and Meloxicam (Boehringer-Ingelheim, Cat#KPOEH3R, *s.c.* 5 mg/kg after surgery every 24 h for 3 consecutive days), *s.c.,* 2 mg/kg after surgery. Animals were sacrificed at P60.

### Transcardiac perfusion

For histological analysis, animals were quickly anesthetized with isoflurane (Zoetis, Cat#6089373) and secured to the perfusion plate as described previously (Venturino et al., 2021). In short: animals were initially perfused with 30 ml of phosphate-buffered saline (PBS) with heparin (100 mg/l, Sigma, Cat#H0878), followed by 30 ml of 4% (w/v) paraformaldehyde (PFA, Sigma, Cat#P6148) in PBS using a peristaltic pump (Behr, Cat#PLP 380, speed: 25 rpm). Animals were decapitated, the brain explanted, fixed in 4% (w/v) PFA for 30 minutes and post-fixed in 4% (w/v) PBS overnight (16h). Then the tissues were washed in PBS and stored at 4°C with 0.025% (w/v) sodium azide (VWR, Cat#786-299). For cryoprotection, the tissue was transferred to 30% (w/v) sucrose (Sigma, Cat#84097) in PBS and incubated overnight at 4°C. To increase antibody permeability, the brain slices were frozen over dry-ice and thawed at room temperature for three cycles.

### Vibratome sections

Cryoprotected samples were embedded into 3% (w/v) Agarose/PBS to obtain coronal brain sections. The brain was sliced in 100 µm coronal sections on a vibratome (Leica VT 1200S).

### Immunofluorescence staining

The brain slices were incubated in blocking solution containing 1% (w/v) bovine serum albumin (Sigma, Cat#A9418), 5% (v/v) Triton X-100 (Sigma, Cat#T8787), 0.5% (w/v) sodium azide (VWR, Cat#786-299), and 10% (v/v) serum (either goat, Millipore, Cat#S26, or donkey, Millipore, Cat#S30) for 1 hour at room temperature on a shaker. Afterwards, the samples were immunostained with primary antibodies diluted in antibody solution containing 1% (w/v) bovine serum albumin, 5% (v/v) triton X-100, 0.5% (v/v) sodium azide, 3% (v/v) goat or donkey serum, and incubated for 48 hours on a shaker at room temperature. The following primary antibodies were used: rat α-CD68 (AbD Serotec, Cat#MCA1957, clone FA-11, Lot 1807, 1:250); goat α-Iba1 (Abcam, ab5076, Lot FR3288145-1, 1:250); rabbit anti-Iba1 (GeneTex, Cat#GTX100042, Lot 41556, 1:750). The slices were then washed 3 times with PBS and incubated protected from light for 2 hours at room temperature on a shaker, with the secondary antibodies diluted in antibody solution. The secondary antibodies raised in goat or donkey were purchased from Thermo Fisher Scientific (Alexa Fluor 488, Alexa Fluor 568, Alexa Fluor 647, 1:2000). The slices were washed 3 times with PBS. The nuclei were labeled with Hoechst 33342 (Thermo Fisher Scientific, Cat#H3570, 1:5000) diluted in PBS for 15 minutes. The slices were mounted on microscope glass slides (Assistant, Cat#42406020) with coverslips (Menzel-Glaser #0) using an antifade solution [10% (v/v) mowiol (Sigma, Cat#81381), 26% (v/v) glycerol (Sigma, Cat#G7757), 0.2M tris buffer pH 8, 2.5% (w/v) Dabco (Sigma, Cat#D27802)].

### Confocal microscopy

Images were acquired with a Zeiss LSM880 upright Airy scan or with a Zeiss LSM700 upright using a Plan-Apochromat 40× oil immersion objective N.A. 1.4. 2 × 2 z-stack tail-images were acquired with a resolution of 1024 × 1024 pixels.

### Image processing

Confocal tile images were stitched using the software Imaris Stitcher 9.3.1.v. Then, the confocal images were loaded in Fiji 1.52e (http://imagej.net/Fiji). To remove the background, the rolling ball radius was set to 35 pixels, and images were filtered using a median 3D filter with x, y, z radii set at 3. Image stacks were exported as .tif files, converted to .ims files using the Imaris converter, and imported into Imaris 8.4.2.v. (Bitplane Imaris).

### Quantification of CD68 volume within cells

Surface renderings were generated on microglia and CD68 z-stacks using the surface-rendering module of Imaris 9.2.v Surfaces were generated with the surface detail set to 0.2 µm. To determine the CD68 surface within microglia, the surface-surface coloc plugin was used. This analysis was performed on the entire image. The total ratio of CD68 volume within microglial volume (CD68-to-microglial volume) was calculated per image. To compute the CD68 fold change, the total CD68-to-microglial volume from each condition (sex/time-point) was scaled to the CD68-to-microglial volume ratio from the respective controls. CD68 fold change > 1 means an increase in CD68 volume, CD68 < 1 means a decrease in CD68 volume. CD68 fold-change = 1 denotes no change in CD68 volume.

### Quantification of microglia density and statistical analysis

The spot-function plugin of Imaris 9.2.v was used to count the number of cells, i.e the soma of iba-1 positive microglia within every confocal image. Microglial cell density was estimated as total number of cells obtained in this way, divided by the size of the imaged sample in mm^2^. Sex averages for microglia from each region were compared with two-sided t.test.

### Reconstruction of 3D-traced microglia

After filtering and background subtraction, images were imported in Imaris 9.2.v (Bitplane Imaris). Microglial processes were 3D-traced with the filament-tracing plugin. Since the filament-tracing plugin provides a semi-automated reconstruction, this eliminates the need for a user-blind approach for selecting representative microglia. New starting points were detected when the largest diameter was set to 12 µm and with seeding points of 1µm. Disconnected segments were removed with a filtering smoothness of 0.6 µm. After the tracing, we manually removed cells that were sitting at the border of the image and were only partially traced so that these cells would not be analyzed. The generated skeleton images were converted from .ims format (Imaris) to .swc format ( Stockley, E. W., et al., 1993) by first obtaining the 3D positions (*x*, *y*, *z*) and the diameter of each traced microglial process using the ImarisReader toolbox for MATLAB (https://github.com/PeterBeemiller/ImarisReader) and then exporting for format standardization using the NL Morphology Converter (http://neuroland.org).

### Analysis of morphometric features

Classic morphometric features were calculated from the .swc files using the functions Length (for total process length), N_branch (for number of branches), N_bifs (for number of branching points) and N_tips (for number of terminal points) from L-measure (Scorcioni et al., 2008) (http://cng.gmu.edu:8080/Lm/). Statistical analysis was performed using scipy.stats (v1.6.2) and scikit-posthocs (v0.6.7). These morphometric features were first tested for normality using the Kolmogorov-Smirnov test (scipy.stats.kstest). After determining the non-normal distribution of the features, we performed non-parametric pairwise tests for independence between measurements from two brain regions using the Kruskal-Wallis test (scipy.stats.kruskal). We used Bonferroni-corrected P values, calculated using Dunn’s test via scikit_posthocs.posthoc_dunn (**Supp. Table 2**-**3**).

### Sholl analysis

Sholl curves were calculated from the .swc files using the sholl_crossings function of the NeuroM Python toolkit (https://github.com/BlueBrain/NeuroM). In brief, concentric Sholl spheres centered on the soma of a given traced microglia are constructed with a given step size radius. The number of microglial processes that intersect each Sholl sphere are determined. This step is performed for each traced microglia in the data. From this, Sholl curves of a microglial population are then calculated as the average number of intersections across the population.

### Topological morphology descriptor (TMD)

A topological data analysis algorithm, the TMD, was used to extract topological phenotypes, called persistence barcodes, from 3D morphological structures (https://github.com/BlueBrain/TMD, Kanari et. al., 2018). In brief, the 3D reconstructed microglia is represented as a tree *T* rooted in its soma. The TMD summarizes this tree by calculating *persistence barcodes*, where each bar represents a persistent microglial process with respect to a filtering function, i.e., the radial distance from the soma. Note that the persistence barcode that the TMD associates with *T* under this filtering function is invariant under rotations about the root and rigid translations of *T* in *R*^3^.

Each bar is described by two numbers: the radial distance, *d_i_*, at which a process originates; and the distance, *b_i_*, when it merges with a larger, more persistent process or with the soma. A bar can be equivalently represented as a point (*d_i_*, *b_i_*) in a *persistence diagram*. We could therefore convolve each point in the persistence diagram with a Gaussian kernel and discretize it to generate a matrix of pixel values, encoding the persistence diagram in a vector, called the *persistence image*.

### Average and bootstrapped persistence images

To construct the *average persistence image* of a given condition, all the persistence barcodes of microglia from the same condition are combined before Gaussian convolution and discretization are performed. We also constructed average persistence images by performing first the Gaussian convolution and discretization of individual microglia persistence barcodes before taking the pixel-wise average. This produced qualitatively similar results.

The bootstrapping method subsamples the microglial population within a given condition, thereby introducing variations around the average persistence image. Starting from the population of all microglia from the same condition, called the *starting population* of size *n*, the persistence barcodes of a pre-defined number of unique microglia, called the *bootstrap size*, are combined to calculate the *bootstrapped persistence image*. We iterate this process a pre-defined number of times, *n_samples_*, with replacement to obtain the *bootstrap sample*.

### Subtraction images and TMD distance

The subtraction image is the pixel-wise difference between two given persistence images. From the subtraction image, the TMD distance can be computed as the sum of the absolute pixel-wise difference. For stability of the TMD distance, we refer the reader to Kanari *et al*. (Kanari et al., 2017)

### Hierarchical clustering

Hierarchical clustering allowed us to find similarities between microglia across several conditions. Hierarchical clustering was done on the basis of the average persistence images. Clusters were then identified hierarchically using the average linkage criterion with the TMD distance metric and was implemented using cluster.hierarchy.linkage from SciPy v1.6.2 (https://www.scipy.org). Dendrograms were generated using cluster.hierarchy.dendrogram to visualize the arrangement of the resulting cluster.

### Dimensionality reduction

#### UMAP

A fast, non-linear dimensionality reduction algorithm, UMAP (Mcinnes et al., 2020) (Uniform Manifold Approximation and Projection), was applied to visualize the high-dimensional pixel space of bootstrapped persistence images using a 2D representation while preserving local and global structures in the bootstrap samples (https://github.com/lmcinnes/umap)(Mcinnes et al., 2020). Given a bootstrap sample containing multiple conditions, a TMD distance matrix containing pairwise distances between bootstrapped persistence images in the bootstrap sample is calculated. Principal components are then obtained using a singular value decomposition of the TMD distance matrix. The first 7 principal components, where the elbow in the singular values is located, were used as input to UMAP with n_neighbors = 50, min_dist = 1.0 and spread = 3.0. Note that we have tested for a wide range of parameter values which did not qualitatively change any of the observations we made in the main text (**Supp. Fig. 2E**).

#### tSNE

An alternative dimensionality reduction algorithm is tSNE ^83^ (t-distributed Stochastic Neighbor Embedding, https://github.com/DmitryUlyanov/Multicore-TSNE) which finds a dimensionality-reduced representation where similar points are pulled closer together while dissimilar points are pushed farther apart with high probability. The first 7 principal components were taken as an input to run tSNE with perplexity = 50.

#### Pseudotemporal ordering

The concept of morphological phenotypes as encoded in the persistence images can be likened to transcriptional phenotypes in single-cell RNA sequencing studies. Bootstrapped persistence images, which encapsulate morphological phenotypes of microglial populations from similar conditions, are comparable. Furthermore, it is reasonable to assume that morphological changes in bootstrapped microglial populations from control to disease conditions occur with incremental differences in the persistence images. This conceptual similarity allowed us to use the pseudo-temporal trajectory inference algorithms that are well-used in the single-cell RNA sequencing community to study the morphological progression during microglial development and degeneration.

#### Palantir

Palantir (Setty et al., 2019) uses principles from graph theory and Markov processes to calculate the pseudo-time and the probability of a cell reaching each of the terminal conditions in the sample (https://github.com/dpeerlab/Palantir). First, the principal components of the bootstrapped persistence images were obtained using palantir.utils.run_pca with n_components = 100 and use_hvg = False. The diffusion maps were then calculated from the PCA projections using palantir.utils.run_diffusion_maps with n_components = 10 and knn = 20 which outputs the Palantir pseudo-times. Harmony (Nowotschin et al., 2019) is then used to construct an augmented affinity matrix from the Palantir pseudo-times to connect together the Palantir pseudo-times and construct a trajectory using a force-directed graph (https://github.com/dpeerlab/Harmony).

#### Monocle

To corroborate the Palantir trajectories, an alternative pseudo-temporal trajectory-inference algorithm called Monocle was employed. Monocle (Cao et al., 2019) uses reverse graph embedding which learns a principal graph which approximates a lower-dimensional manifold to construct a pseudo-time trajectory (https://github.com/cole-trapnell-lab/monocle3)(Cao et al., 2019). Similar to Palantir implementation, the principal components of the bootstrapped persistence images were first obtained using preprocess_cds with num_dim = 100. A 2D UMAP representation was then obtained using reduce_dimension with umap.metric = “manhattan”, umap.min_dist = 1.0, and clusters were identified using cluster_cells with cluster_method = ’leiden’. Finally, the pseudo-temporal trajectory was then obtained using learn_graph with use_partition = FALSE and close_loop = FALSE.

#### Stable ranks analysis

An alternative representation of the persistence barcodes is through stable ranks (Riihimäki and Chacholski, 2018). Stable ranks are functional summaries of persistence which depend on pseudometrics to compare persistence barcodes. Given a pseudometric *d*, the stable rank 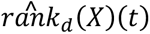 of a persistence barcode *X* is a function that assigns to *t* the number:

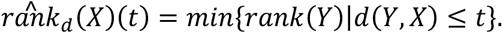

whereby *rank(Y)* denotes the number of bars of the persistence barcode *Y*. The stable rank 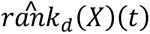 associates to a persistence barcode a non-increasing and piece-wise constant function with values in [0, ∞). An important property is that this mapping is continuous with respect to the chosen pseudometric *d* and the *L_p_* metric on the space ℳ of measurable functions.

A class of pseudometrics on persistence barcodes can be constructed from density functions (Riihimäki and Chacholski, 2018), which intuitively are used to vary the weight along the filtration scale parametrizing a barcode. With such pseudometrics, the stable rank is a bar count based on length of bars as scaled by the density. The standard stable rank is defined by a density function with constant value one. In this case, 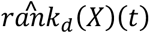 is the number of bars in *X* with length greater than or equal to *t*, i.e., all filtration scales are weighted equally.

Stable ranks can be used in place of persistence images in the MorphOMICs pipeline. Similarly, to MorphOMICs, the persistence barcode *X* of a given microglia is calculated using the TMD algorithm. To obtain *bootstrapped standard stable ranks*, we combined the persistence barcodes of a pre-defined number of microglia and computed their standard stable ranks. Dimensionality reduction was then implemented similar to the methods above (see **Methods: *Dimensionality reduction***).

#### Classification accuracy using stable ranks

To support and quantify the impact of bootstrapping on the regional segregation visualized in the reduced UMAP space (**Fig. 1F**), we performed a classification task for microglia morphologies represented by their standard stable rank and labeled by brain region. We used a support vector machine (SVM) with a specific kernel based on stable ranks (Agerberg et al., 2021) for the classification. For persistence barcodes *X* and *Y*, the stable rank kernel with respect to a pseudometric *d* is given by

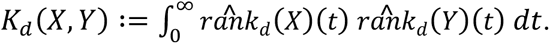

where we used the pseudometric induced by the constant function with value one.

We performed pairwise classifications. For each pair of brain regions, we constructed a dataset consisting of 400 bootstrap samples, i.e., 200 from each region and bootstrap sizes of either 10, 20 or 50 (the results are reported separately for these three values). We randomly partition the dataset for cross-validation wherein 240 samples were used for SVM training (training set) and 160 samples for validation (test set). We report the average accuracy over 10 repeated cross-validations on the test set. The SVM was trained using the implementation in the Python library sklearn (https://scikit-learn.org/stable/) with default settings except for the usage of the stable rank kernel.

#### Bootstrapped morphometric features and bootstrapped Sholl curves

To understand whether classical morphology analysis pipelines are able to recapitulate the microglial dynamics recovered by MorphOMICs, a similar bootstrapping analysis was also done where we pooled together a pre-defined number of microglia. Each morphometric quantity in the extended list enumerated in **Supp. Table 4** was then averaged to obtain a 27-D vector, with each dimension corresponds to a morphometric feature, called the *bootstrapped morphometric features*. On the other hand, Sholl curves averaged across the pooled microglia to obtain the *bootstrapped Sholl curves*. Dimensionality reduction was then implemented similar to the methods above (see **Methods: *Dimensionality reduction***).

### Data availability

All the .swc files are available upon request.

## Acknowledgements

We thank the scientific service units at IST Austria, in particular Michael Schunn’s team at the preclinical facility, and especially our colony manager Sonja Haslinger, for excellent support. We are also grateful to the IST bioimaging facility, and in particular Christoph Sommer for helping with the data file conversions. We thank Margaret Maes, Balint Nagy, Sara Oakeley, Marco Benevento and all members of the Siegert group for constant feedback on the project and on the manuscript. This research was supported by the European Union Horizon 2020 research and innovation program under the Marie Skłodowska-Curie Actions program (754411 to R.J.A.C.), and by the European Research Council (grant No. 715571 to S.S.). L.K. was supported by funding to the Blue Brain Project, a research center of the École polytechnique fédérale de Lausanne, from the Swiss government’s ETH Board of the Swiss Federal Institutes of Technology.

## Author contributions

Conceptualization, G.C., R.J.A.C, L.K., M.S., J.A., W.C., K.H., S.S.; Methodology, G.C., R.J.A.C., L.K., S.S.; Software, R.J.A.C., L.K.; Validation, G.C., R.J.A.C.; Formal analysis, G.C., R.J.A.C., L.K.; Investigation, G.C., R.J.A.C., A.V., R.S., S.S.; Resources, L.K., H.M., L-H.T.; Data Curation, G.C., R.J.A.C.; Writing – Original Draft and Visualization, G.C., R.J.A.C., S.S.; Supervision, Project Administration, Funding Acquisition, S.S.; Stable ranks: Conceptualization, G.C., R.J.A.C., L.K., M.S., J.A., W.C., K.H., S.S.; Software Validation, and Formal Analysis, R.J.A.C., J.A.

## Computational assessment of bootstrapping methods in MorphOMICs

### 1. Single-condition case

Microglia are highly dynamic (Tremblay et. al., 2011) – a feature which is inherent to their function. Under homeostatic conditions, microglia survey their local environment for insults and anomalies. This intrinsic variability challenges the topological analysis of microglial morphology as we observed in the corresponding persistence images of single microglia (**Fig. 1B**). This variability can mask heterogeneity between microglial populations from different biological conditions *e.g.*, brain region, sex, and development and disease time point (**Supp. Fig. 2B**).

To overcome this intrinsic variability within microglial populations, we use bootstrapping methods. Bootstrapping is a statistical method that combines random resampling and permutation. It is commonly used to calculate standard errors, to construct confidence intervals, and to perform hypothesis testing for numerous types of sample statistics. In our case, we used bootstrapping to randomly pool together a pre-determined number of microglia within a condition, called the bootstrap size, to reduce the dispersion. This allowed us to construct a bootstrapped persistence image of this microglial sub-population. By averaging out the highly variable portions of the persistence images, we retain the topological signatures that may separate different conditions. Moreover, bootstrapping makes it possible to create as many bootstrapped persistence images as desired.

The pixels of the persistence image span a high-dimensional space. The bootstrapped persistence images form a point cloud in this high-dimensional space with the average persistence image in the center (**Supp. Fig. 2A**). Thus, the spread of this cloud allows us to assess the variability within a microglial population. Intuitively, the bootstrap size affects this point cloud size. To construct the bootstrapped persistence images with just a single microglia will give us the largest cloud as it reflects the full size of the population’s dispersion. On the other hand, when the bootstrapped persistence image is constructed using all of the microglia, the cloud collapses to a single point as there is no difference between the bootstrapped persistence images and the average persistence image.

To systematically understand the effect of the bootstrap size on the structure of the point cloud formed by the bootstrapped persistence images, we considered a population composed of a high number of traced microglia, namely the microglia of the adult, healthy dentate gyrus. This allowed us to span a range of sizes for the starting population from which we performed the bootstrapping. First, we only considered microglia traced from male animals to avoid inter-sex differences (**Fig. 1F**). As we observed no animal-specific batch effects (**Supp. Fig. 1C**), we selected four mice with the highest number of tracings, and grouped them into two artificial groups (A and B) so that the number of single cells were similar in each group (*N_A_* = 223, *N_B_* = 231). Despite coming from the same brain region, these two groups have a non-zero TMD distance (*d* = 10.64) which we call the TMD intrinsic distance. This intrinsic distance arises due to small but accumulated variations in the persistence images. To account for the effect of unequal starting population sizes between the groups, we randomly selected *n* = 200 traced microglia from each group to form the starting population. We then drew single microglia from these groups to create a set of bootstrapped persistence images, which we call bootstrap samples A and B. Note that the TMD intrinsic distance remains the same, regardless of the bootstrap size (**Supp. Fig. 2A**).

To characterize the point cloud formed by the bootstrapped persistence images, we calculated the within-condition distance which is the average TMD distance between two persistence images within the same bootstrap sample. We want to stress here that the two conditions A and B are artificial: the bootstrap persistence images in bootstrap samples A and B come from the same brain region. We observed that reducing the bootstrap size compacted the point cloud and subsequently reduced the within-condition distance within the bootstrap sample (**Supp. Fig. 2C**). At a certain bootstrap-to-starting population size ratio where *N_A_* = *N_B_* = 200 the within-condition distance becomes smaller than the TMD intrinsic distance. This implies that the between-condition distances, *i.e.* the TMD distance between persistence images across different bootstrap samples, increases causing a forced separation between two groups from the same condition.

Thus, it is imperative to select a bootstrap size which reduces the dispersion of the bootstrap samples without artificially separating samples that share topological signatures. One way to address the latter condition is to determine whether the bootstrap samples A and B cluster separately under a given bootstrap size. To test this, we performed a complete-linkage hierarchical clustering with the TMD distance as the metric. We imposed a cut-off which results in two clusters, *ω*_1_ and *ω*_2_. We then defined a mixing entropy of the resulting clusters Ω = { *ω*_1_, *ω*_2_} which measures the discrimination between bootstrap samples and is calculated as

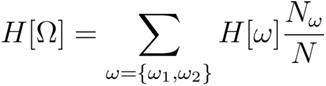

where 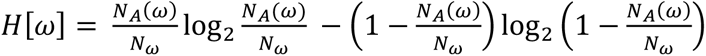 is the entropy of cluster*ω*, *N_A_(ω)* is the number of bootstrapped persistence images in bootstrap sample A located in cluster *ω* and 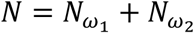 is the total number of bootstrapped persistence images in the point cloud. To understand this mixing entropy, we considered both extreme situations. The first case is when bootstrap samples A and B overlap and the point cloud is highly dispersed: this results in a big cluster with all persistence images aggregated except for one. The latter is the last bootstrap persistence image to be clustered in the dendrogram, and the mixing entropy is close to 1. On the other hand, when the point cloud dispersion was small enough for bootstrap samples A and B to segregate, the clustering resulted in separate clusters for samples A and B, and the mixing entropy is zero. Note, the latter emerges whenever we force the separation between two similar-condition groups.

If we also consider the situation between the two extreme situations, the mixing entropy decreases with increasing bootstrap-to-starting population size ratio (**Supp. Fig. 2C**). We observed that the mixing entropy remained close to 1 for a range of small ratios and then dropped to zero, depending on the starting population size. This behavior allows us to define an optimal bootstrap size which maximizes the trade-off between the intrinsic variability of the bootstrap samples and the indistinguishability of samples coming from the same conditions. By dividing the mixing entropy with the within-condition distance, we found a peak close to a bootstrap size that is 30% of the starting population size (**Supp. Fig. 2D**).

Furthermore, we assessed the effect of low starting population sizes on both the within-condition distances and mixing entropy. Thus, we randomly selected *n* = {10, 20, 30, 40, 50, 60, 70, 80} microglia from each group and performed the bootstrapping over the resulting starting populations. We found that both within-condition distance and mixing entropy decreased as a function of the bootstrap size, but dependent on the starting population size (**Supp. Fig. 2C**). Interestingly, when we divided the mixing entropy by the within-condition distance, we observed that there was no longer a well-defined optimal bootstrap size (**Supp. Fig. 2D**), and that the optimal ratio is a range of parameters which is larger for samples with a low starting population size.

### 2. Multiple-condition case: sexual dimorphism in the frontal cortex and dentate gyrus

In the analyses above, we only considered the situation where two groups come from the same condition, such as microglia from the adult male dentate gyrus. Here, we look at a situation where we have multiple conditions which not only exhibit spatial but also sexual heterogeneity.

We focused on microglial populations coming from the healthy, adult frontal cortex (FC) and the dentate gyrus (DG) where we see a microglial signature and a region-dependent sexual dimorphism. Thus, we took 75 microglia from the four male and four female mice with the highest number of traced microglia in the frontal cortex and dentate gyrus. To investigate the effect of having unequal proportions of male and female microglia in the sample, we created starting populations with size *N* = 200 where the male-to-female microglia ratio, *r* = {0.0, 0.1, 0.2, … , 0.8, 0.9, 1.0} was fixed. For each brain region and male-to-female ratio, we constructed bootstrap samples with bootstrap-to-starting population size ratio at 0.1, 0.3, 0.5, 0.7, and 0.9. As there are multiple conditions, we looked at the 2D UMAP representations for each bootstrap size using the same parameters in the main text.

We observed that at 0.1 size ratio, the difference in the morphological signature between FC and DG microglia is already apparent (**Supp. Fig. 2H**), and that within a brain region cluster, the pure male and female samples are located at opposite ends. These ends in the DG_mg_ cluster tend to come closer together than those of the FC_mg_. Note that these observations are captured when the bootstrap-to-starting population size ratio is at 0.3.

Finally, we observed that as the bootstrap size increased, samples with different male-to-female size ratios broke apart, suggesting a forced separation between different conditions. However, the rate at which the different conditions became more distinct was not uniform. Indeed, we saw that the male-dominated samples in the DG_mg_ still form a cluster at bootstrap-to-starting population size ratio 0.9, which implies that ♂DG_mg_ have “stronger” and less variable microglial signatures than their female counterparts. This suggests that spanning a range of size ratios can uncover information on the intrinsic variability.

